# A minimal CRISPR polymerase produces decoy cyclic nucleotides to detect phage anti-defense proteins

**DOI:** 10.1101/2025.03.28.646047

**Authors:** Ashley E. Sullivan, Ali Nabhani, Kate Schinkel, David M. Dinh, Melissa L. Duncan, Eirene Marie Q. Ednacot, Charlotte R.K. Hoffman, Daniel S. Izrailevsky, Emily M. Kibby, Toni A. Nagy, Christy M. Nguyen, Uday Tak, A. Maxwell Burroughs, L. Aravind, Aaron T. Whiteley, Benjamin R. Morehouse

## Abstract

Bacteria use antiphage systems to combat phages, their ubiquitous competitors, and evolve new defenses through repeated reshuffling of basic functional units into novel reformulations. A common theme is generating a nucleotide-derived second messenger in response to phage that activates an effector protein to halt virion production. Phages respond with counter-defenses that deplete these second messengers, leading to an escalating arms race with the host. Here we discover a novel antiphage system we call Panoptes that detects phage infection by surveying the cytosol for phage proteins that antagonize the nucleotide-derived second messenger pool. Panoptes is a two-gene operon, *optSE*. OptS is predicted to synthesize a second messenger using a minimal CRISPR polymerase (mCpol) domain, a version of the polymerase domain found in Type III CRISPR systems (Cas10) that is distantly related to GGDEF and Thg1 tRNA repair polymerase domains. OptE is predicted to be a transmembrane effector protein that binds cyclic nucleotides. *optSE* potently restricted phage replication but mutant phages that had loss-of-function mutations in anti-CBASS protein 2 (Acb2) escaped defense. These findings were unexpected because Acb2 is a nucleotide “sponge” that antagonizes second messenger signaling. Using genetic and biochemical assays, we found that Acb2 bound the OptS-synthesized nucleotide, 2′,3′-cyclic adenosine monophosphate (2′,3′-c-di-AMP); however, 2′,3′-c-di-AMP was synthesized constitutively by OptS and inhibited OptE. Nucleotide depletion by Acb2 released OptE toxicity thereby initiating abortive infection to halt phage replication. These data demonstrate a sophisticated immune strategy that hosts use to guard their second messenger pool and turn immune evasion against the virus.

## Introduction

Bacteria are under constant assault from their viruses, bacteriophages (phages), and have evolved sophisticated mechanisms to protect themselves^1,2^. The immune system of any bacterium is a combination of individual pathways called antiphage systems that limit phage replication. Bacteria encode an average of 5.8 antiphage systems per strain^3^. However, each bacterial strain within a species dynamically exchanges antiphage systems through horizontal gene transfer giving each a different suite of defenses^1^.

Antiphage systems must rapidly sense phage infection and transduce an activation signal to an effector protein to stop virion production. Recent progress cataloging antiphage systems has identified that many systems transduce their activation signal through nucleotide-derived second messengers. The cyclic oligonucleotide-based antiphage signaling system (CBASS) synthesizes cyclic di– and trinucleotides^4–6^, the pyrimidine cyclase system for antiphage resistance (Pycsar) synthesizes cyclic mononucleotides^7^, and the Thoeris system synthesizes cyclic ADP-ribose derivatives in response to phage^8^. The advantage of nucleotide-derived second messengers is they can be synthesized from abundant precursors and amplify a rare phage-sensing event into stoichiometrically more activated effector proteins. The success of these strategies is further underscored by their remarkable spread through lateral transfer: homologs of CBASS components can be found in the cGAS-STING pathway of metazoans and homologs of Thoeris components can be found in the immune system of plants and metazoans^9,10^.

Nucleotide-derived second messengers are also a vulnerability because viruses use phosphodiesterases and “sponge proteins” to interrupt signaling^11–18^. Just as the nucleotide second messenger strategy is pervasive from bacteria to eukaryotes, so too are some of the immune evasion strategies from viruses that infect bacteria and metazoans^19,20^. Bacterial antiphage systems synthesize a wide variety of chemical variations of their class of second messenger, often using different nucleotide bases, forming different phosphodiester linkages, or even incorporating amino acids^6,7,21^. This variability is likely the result of the arms race between host and pathogen. Nevertheless, most viral proteins that antagonize second messengers can interact with a wide range of second messenger variants suggesting that changing the second messenger is not sufficient to out-pace the virus^11–14,17,18,22–24^.

Here we investigated a two-gene operon (*optSE*) that appeared similar to CBASS, Pycsar, and Thoeris systems with a predicted nucleotide second messenger synthase (OptS) and a nucleotide-binding effector protein (OptE). However, unlike systems that synthesize signaling nucleotides in response to phage, we found that OptS constitutively produces a signaling nucleotide to repress growth inhibition by OptE. The OptS nucleotide, 2′,3′-c-di-AMP, is similar to nucleotides found in CBASS systems, which is synthesized by a SMODS domain containing protein called a CD-NTase. Phages that encoded anti-defense proteins that deplete cyclic nucleotides to circumvent CBASS signaling inadvertently depleted OptS-synthesized 2′,3′-c-di-AMP and unmasked OptE toxicity, which interrupted the phage replication cycle through destruction of the host cell, a process called abortive infection. In this way, OptSE guards the cyclic dinucleotide pool of the cell and detects phage infection by sensing immune evasion. We named OptSE the Panoptes antiphage system for Argus Panoptes, the all-seeing, many-eyed giant in Greek mythology who was a faithful watchman to Hera.

## Results

### The Panoptes system is antiphage

We investigated a candidate two-gene Panoptes operon from *Vibrio navarrensis*. The first gene (*optS*, S = <u>S</u>ynthase) encoded a minimal CRISPR polymerase (mCpol) domain that is predicted to synthesize a nucleotide second messenger^25^. The second gene (*optE*, E = <u>E</u>ffector) encoded two predicted transmembrane domains and a SMODS-associating 2TM, β-strand rich (S-2TMβ) domain (**Fig. 1a**)^25^. OptE is homologous to Cap15 proteins within CBASS that disrupt membrane integrity in response to nucleotide^25,26^. OptSE resembled CBASS^5^, Pycsar^7^, and Thoeris^8^ systems with a nucleotide-derived second messenger-generating protein and a second messenger-responsive effector protein. We expressed the operon from its endogenous promoter on a medium-copy plasmid in *E. coli* MG1655 and challenged these bacteria with a panel of diverse phages. The *optSE* operon specifically defended against phages from the Straboviridae (**Fig. 1b)**. We focused on the *Tequatroviruses* genus that includes the T-even phages and found *optSE* provided over 100-fold protection against phage BASEL 38, 41, 42, and 44, and over 500-fold protection for BASEL36, 37, 39, 40, and 43 (**Fig. 1b and c**).

**Figure 1.**
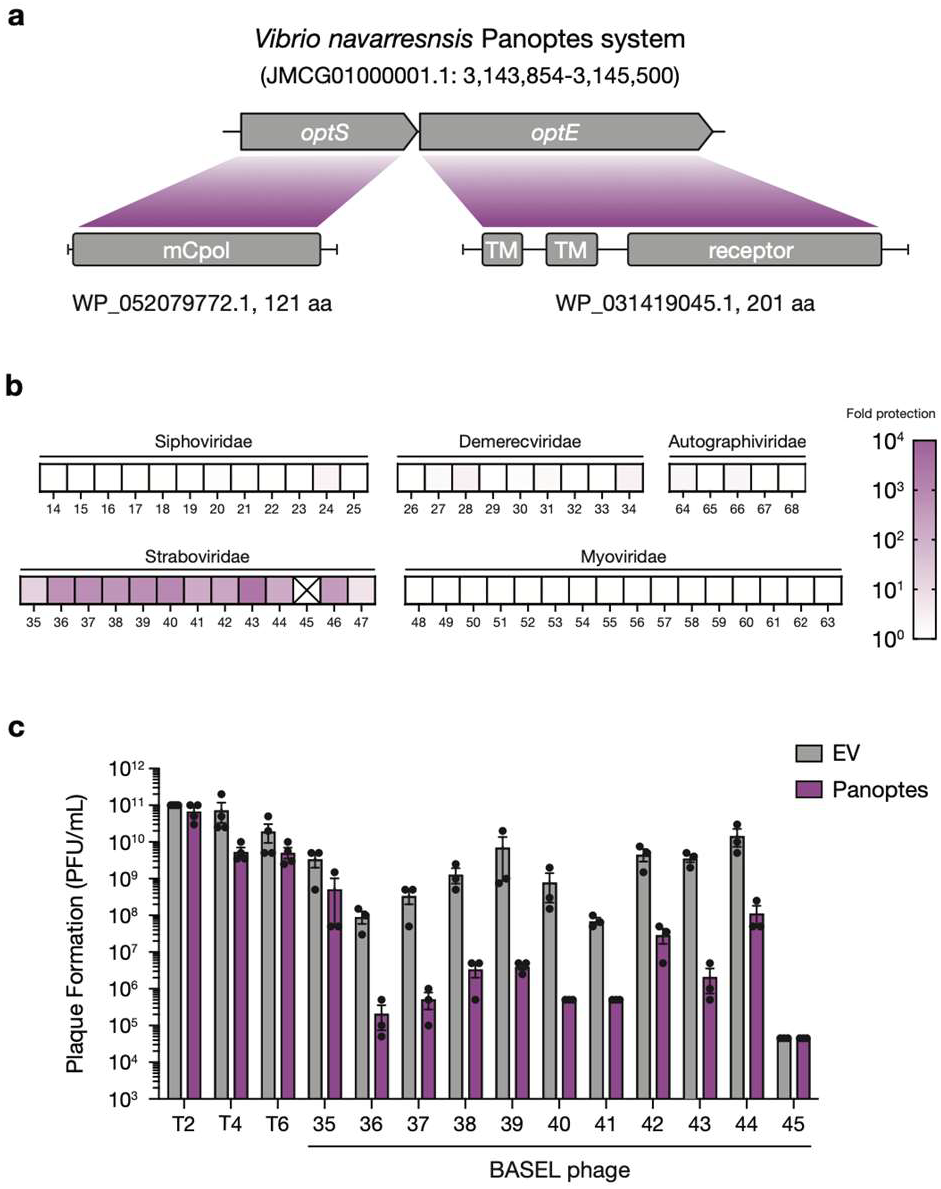
The Panoptes system is antiphage. **(A)** Domain architectures of Panoptes operon genes, *optS* and *optE*. **(B)** Heatmap of fold defense provided by the Panoptes system for a panel of diverse phages from the BASEL collection. *E. coli* expressing the Panoptes system were challenged with phages and fold defense was calculated for each phage by dividing the efficiency of plating (in PFU/mL) on empty vector by the efficiency of plating on Panoptes system-expressing bacteria. An “X” represents uncountable plaques. Family names are above each indicated set of phages. **(C)** Efficiency of plating of indicated phages infecting *E. coli* expressing a plasmid with either the Panoptes system or an empty vector. Data represent the mean ± standard error of the mean (SEM) of n = 3 biological replicates, shown as individual points.

We selected the best characterized T-even phage, phage T4, for additional characterization. We confirmed that *optSE* provided defense against phage T4 in both soft-agar and liquid cultures (**Fig. 1c**; **ED Fig. 1**). A somewhat surprising outlier was phage T2, which was not restricted by *optSE*. These findings were unexpected as phage T2 is generally much more sensitive to phage defense systems than phage T4^27,28^. Taken together, the results support that Panoptes is an antiphage system.

### OptS is a minimal CRISPR polymerase

To better understand the possible enzymatic function of the OptS protein, we expressed and purified an ortholog from *K. pneumoniae* KP67 (*Kp*OptS) for *in vitro* biochemical and structural characterization experiments. While *Kp*OptS is a small protein (∼13 kDa) based on primary amino acid sequence alone, size exclusion chromatography supported the existence of a stable tetrameric species in solution (**Fig. 2a**). We next determined the crystal structure of *Kp*OptS, which could be refined at a nominal resolution of 1.75 Å (**ED Table 1, See methods**). The crystallographic asymmetric unit contained four copies of *Kp*OptS, forming an apparent tetrameric complex, which further supports that *Kp*OptS likely exists as a tetramer (**Fig. 2b; ED Fig. 2a-c**). Minimal conformational and B-factor variation was observed across the protomers and superposition revealed RMSD values of ∼0.3-0.4 Å on average between pairs.

**Figure 2.**
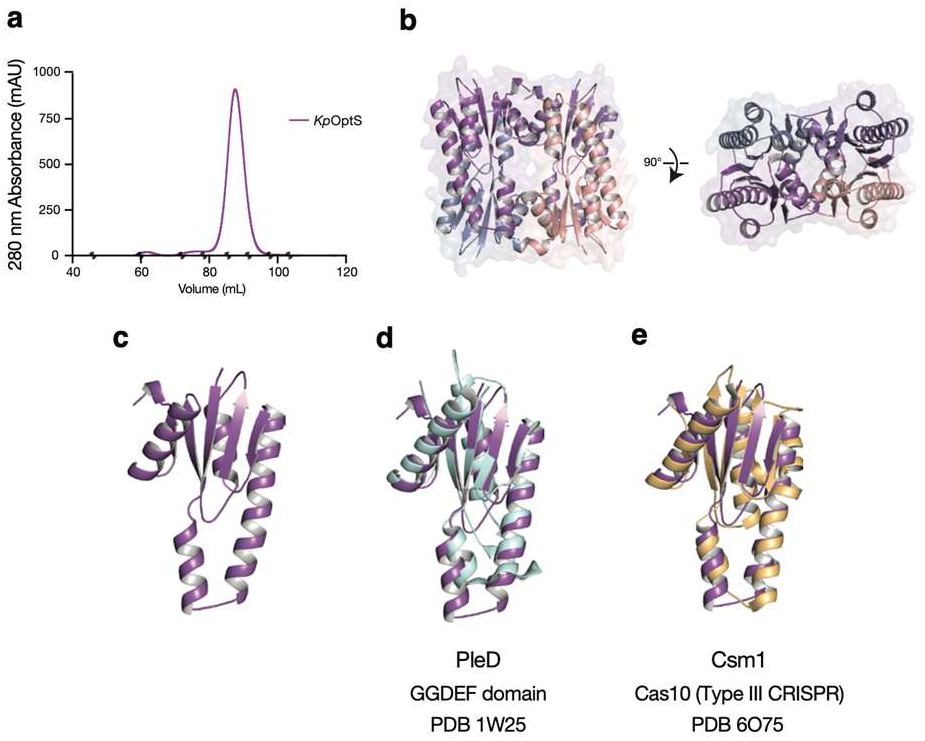
OptS is a minimal CRISPR polymerase. **(A)** Size exclusion chromatography reveals a monodisperse signal for *Kp*OptS with an elution volume consistent with an oligomer of at least a trimer. Grey dots indicate molecular weight standards: Blue dextran, 46.2 mL, 2000 kDa; Ferritin, 59.8 mL, 440 kDa; Aldolase, 71.0 mL, 158 kDa; Conalbumin, 78.9 mL, 75 kDa; Ovalbumin, 85.3 mL, 43 kDa; Carbonic anhydrase, 91.5 mL, 29 kDa; Ribonuclease A, 98.6 mL, 13.7 kDa; Aprotinin, 104.4 mL, 6.5 kDa. **(B)** Crystal structure of the *Kp*OptS apoprotein reveals a tetrameric architecture. Two rotated views are depicted with each protomer shown in a different color. **(C)** Structure of the isolated protomer (monomer) highlighting overall domain fold. **(D)** Superposition of GGDEF-containing enzyme PleD (PDB 1W25, light blue) with *Kp*OptS. **(E)** Superposition of Type III Cas10 CRISPR polymerase palm domain (PDB 6O75, light orange) with *Kp*OptS.

The protomer of *Kp*OptS contains a <u>R</u>NA-recognition <u>m</u>otif (RRM, sometimes referred to as ferredoxin) polymerase palm-domain fold composed of four primary α-helices surrounding a five-stranded β-sheet (**Fig. 2c**)^25^. Although quite different in quaternary structure, the protomeric unit of *Kp*OptS aligns well with the palm domains of the Cas10 (Csm1) from *Thermococcus onnurineus* and the GGDEF diguanylate cyclase PleD from *Caulobacter crescentus* with RMSD values of 3.86 Å and 2.39 Å, respectively (**Fig. 2d and e**)^29,30^. These polymerase domains were also earlier unified with the histidinyl tRNA repair polymerase Thg1^31^. Indeed, the OptS fold observed in the crystal structure was previously predicted by Burroughs et al. and described as a minimal CRISPR polymerase (mCpol) owing to predicted homology to Cas10 enzymes from type III CRISPR immune systems^25^. GGDEF diguanylate cyclases synthesize cyclic diguanosine monophosphate (c-di-GMP) from GTP precursors and Cas10 enzymes synthesize cyclic oligoadenylate molecules (between 2–6 AMPs) from ATP precursors^32–34^. The high degree of structural similarity to GGDEF, Cas10, and other nucleotide polymerases (palm domains are found in a variety of nucleotidyltransferases such as the SMODS of CBASS and cUMP/cCMP synthases of Pycsar defense systems) strongly implicates Panoptes OptS proteins in the enzymatic processing of nucleotides.

### OptS synthesizes 2′,3′-c-di-AMP

Purified *Kp*OptS was incubated with ribonucleotide triphosphates (ATP, GTP, CTP, UTP) and the reactions were separated and monitored by high-performance liquid chromatography (HPLC) to detect any synthesis activity (**ED Fig. 3a-d**). The largest abundance product observed was derived from ATP with minor products requiring the coincubation of both ATP and GTP (**Fig. 3a; ED Fig. 3b and c**). We sought to determine the identity of the ATP-derived *Kp*OptS product and found that it had a similar retention time to an isomer of cyclic diadenosine monophosphate with mixed phosphodiester linkages, 2′,3′-c-di-AMP (c[A(2′– 5′)pA(3′–5′)p]) (**Fig. 3b**). Additional biochemical evidence for both the cyclic and mixed phosphodiester linkages of the product came from nuclease treatment experiments (**ED Fig. 3e**). The product was resistant to calf-intestinal phosphatase (CIP, cleaves terminal phosphates) but was partially susceptible to P1 nuclease (cleaves 3′–5′ phosphodiester bonds) shifting to a new retention time. The combination CIP and P1 treatment led to loss of signal at retention times observed in the untreated and P1-alone treatment conditions and the appearance of a new, late eluting, peak that was consistent with A(2′–5′)pA generated by CIP treatment of commercially available linear pppA(2′–5′)pA.

**Figure 3.**
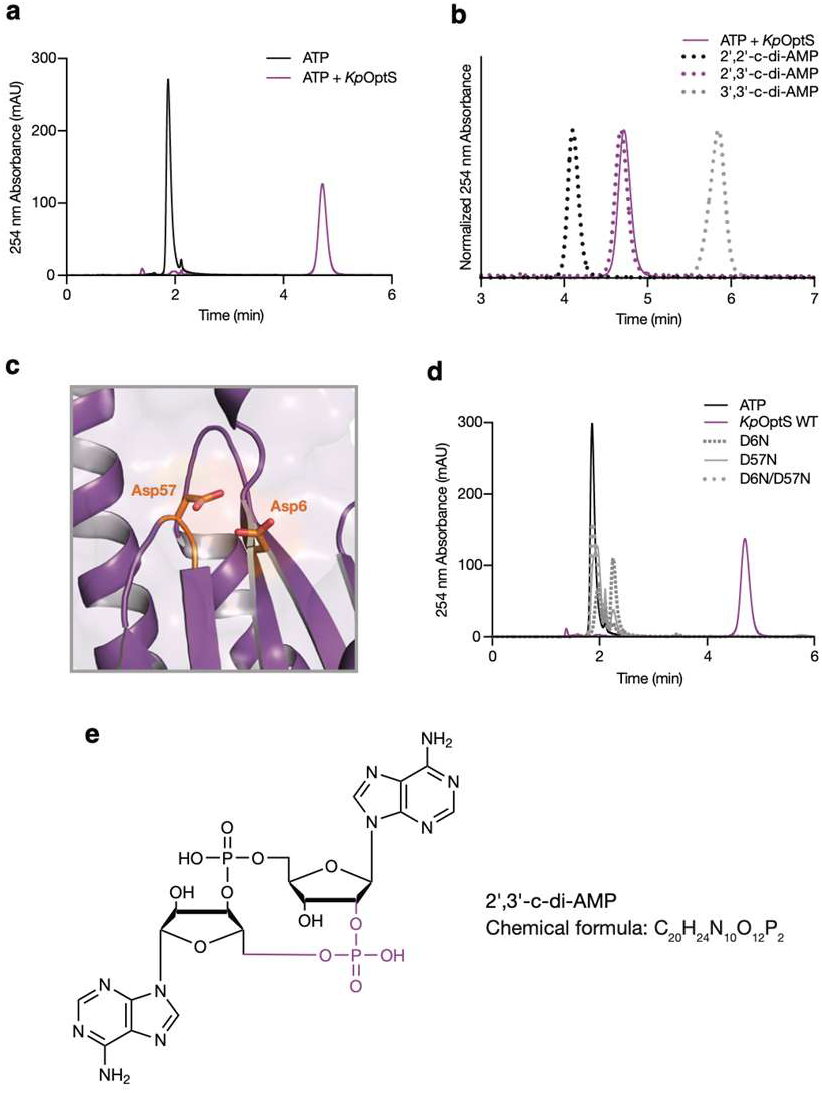
OptS is capable of synthesizing 2′,3′-c-di-AMP. **(A)** HPLC analysis of an ATP chemical standard compared with the product of *Kp*OptS when incubated with ATP alone. Traces representative of at least three replicates. **(B)** HPLC analysis of 2′,2′-c-di-AMP, 2′,3′-c-di-AMP, and 3′,3′-c-di-AMP chemical standards compared with the product of *Kp*OptS. Traces representative of at least three replicates. **(C)** Magnified view of the *Kp*OptS putative active site highlighting catalytic Asp6 and Asp57 (orange). **(D)** HPLC analysis of the reaction products when wild type or mutant *Kp*OptS is incubated with ATP, compared to an ATP chemical standard. Traces representative of at least three replicates. **(E)** Chemical structure of 2′,3′-cyclic di-adenosine monophosphate (2′,3′-c-di-AMP).

Based on conservation with diverse nucleotidyltransferases, we expected that OptS Asp6 and Asp57 would be involved in catalysis as they likely coordinate Mg^2+^ ions for triphosphate stabilization in the binding pocket (**Fig. 3c; ED Fig. 3f**). Accordingly, mutation of these amino acids (either to Ala or Asn) alone or in tandem led to complete loss of cyclic dinucleotide production (**Fig. 3d**). Finally, liquid chromatography-mass spectrometry (LC-MS) analysis of the *Kp*OptS reaction with ATP confirmed that the main product is indeed 2′,3′-c-di-AMP (**Fig. 3e; ED Fig. 3g**).

### Acb2 is necessary for activation of the Panoptes system

Phage T4 formed irregular plaques in soft-agar overlays of *optSE*-expressing bacteria (**ED Fig. 4a**). At higher concentrations of phage, large clear plaques appeared and we hypothesized that these large plaques might be caused by mutant phages capable of escaping *optSE*-mediated defense. We isolated escaper mutants from three unrelated, clonal T4 lysates. 15 candidate escapers were isolated, plaque purified, and genome sequenced along with their parent wild-type phages (**ED Fig. 4b**). Surprisingly, 14 of 15 escaper phages encoded mutations exclusively in the anti-CBASS 2 gene (*acb2/vs*.4; **Fig. 4a; ED Table 2**). Mutations included amino acid substitutions (E26K, C78W, F82C, or P84Q), insertions, and deletions that caused frame shifts, premature stop codons, and large-scale deletions.

**Figure 4.**
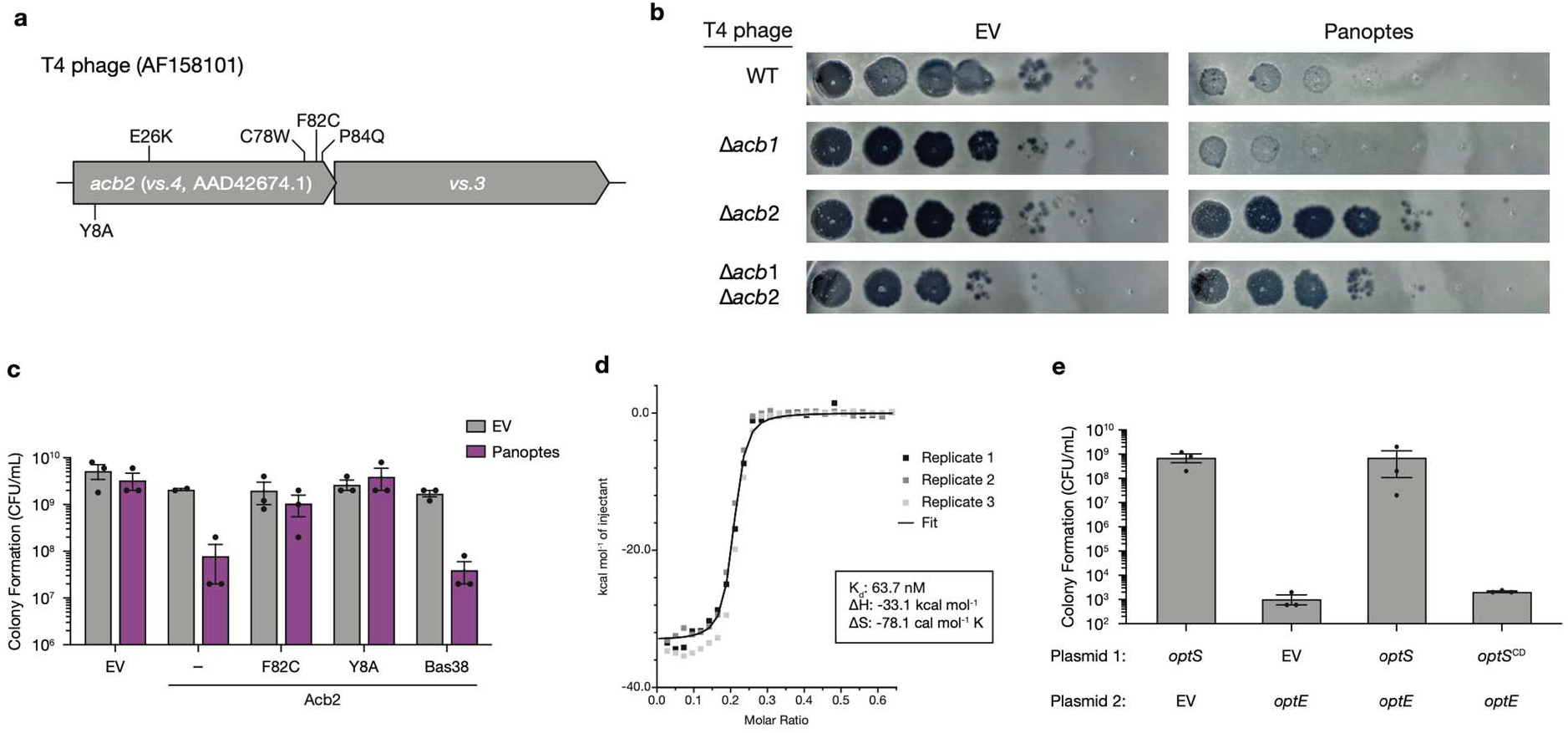
Acb2 is necessary and sufficient for activating the Panoptes system. **(A)** Schematic of phage T4 escape mutations from this paper (top) and a previously validated binding site mutation (bottom). **(B)** Efficiency of plating of T4 phages with the indicated genotype on *E. coli* expressing an empty vector or the Panoptes operon from a plasmid. Images are representative of n=3 biological replicates. **(C)** Colony formation of *E. coli* expressing an empty vector or Panoptes on a low copy plasmid and indicated empty vector or Acb2 allele on an IPTG-inducible plasmid. Data represent the mean ± SEM of n = 3 biological replicates, shown as individual points. **(D)** Isothermal titration calorimetry (ITC) to test binding of 2′,3′-c-di-AMP to Acb2. The individual data points of three technical replicates are shown. The K_d_, ΔH, and ΔS were determined by generating a global fit for the three indicated replicates. Raw data for these curves are shown in Extended Data Figure 5. **(E)** Colony formation of *E. coli* co-expressing indicated genes from IPTG-inducible (Plasmid 1) and arabinose-inducible (Plasmid 2) plasmids. Data represent the mean ± SEM of n = 3 biological replicates, shown as individual points.

Acb2 was recently discovered as an anti-defense protein that antagonizes antiphage immunity by acting as a nucleotide “sponge,” sequestering CBASS-derived cyclic oligonucleotide signaling molecules^12,13,22^. Acb2 binding of CBASS-derived nucleotides interrupts downstream effector protein activation, allowing the phage to evade defense. We were surprised to find that escaper phages harbored obvious loss-of-function mutations in Acb2 because this ruled out that our escapers had simply mutated Acb2 to increase affinity for the OptS-derived nucleotide. Instead, these data suggested that Acb2 was necessary for OptSE-mediated defense.

We tested whether Acb2 was necessary for OptSE-mediated defense by challenging empty vector (EV) or *optSE*-expressing bacteria with either wild-type or Δ*acb2* T4 phages. Wild-type T4 efficiently replicated on EV-expressing bacteria and was restricted by *optSE*. T4 Δ*acb2*, however, was no longer restricted by OptSE and similarly replicated in both the EV- and *optSE*-expressing bacteria (**Fig. 4b**). Phage T4 also encodes anti-CBASS 1 (*acb1*), a phosphodiesterase that degrades cyclic oligonucleotides to similarly antagonize CBASS signaling^11^. We constructed a T4Δ*acb1* phage and found no impact on *optSE*-mediated defense (**Fig. 4b; ED Fig 4c**). Nevertheless, the T4Δ*acb1*Δ*acb2* double mutant appeared similar to T4 Δ*acb2* alone. Taken together, these data suggest that the phage gene *acb2*, but not *acb1*, is necessary for *optSE*-mediated defense.

### Acb2 is sufficient for activation of the Panoptes system

We next explored whether *acb2* is sufficient to activate Panoptes in the absence of phage. OptE is a member of the S-2TMβ family of proteins that are comprised of a two-pass transmembrane element obligately fused to a soluble β-barrel nucleotide-binding domain. Another member of this family, Cap15, is a CBASS effector that disrupts the bacterial inner membrane to initiate abortive infection^26^. We hypothesized that OptE might similarly disrupt bacterial host membranes and inhibit colony formation. Therefore, we co-expressed the *optSE* operon with *acb2* and assayed for colony formation. T4 *acb2* selectively inhibited colony formation only when co-expressed with the *optSE* operon (**Fig. 4c**). We observed the same phenotype when we expressed *acb2* from phage BASEL 38, which is restricted by the *optSE* operon by over 400-fold. These data show that Acb2 is both necessary and sufficient to activate OptSE.

Using the same co-expression assay, we found that the mutant *acb2* allele encoding Acb2^F82C^ identified in our escaper phage did not inhibit colony formation. The structure of Acb2 from phage T4 shows the F82 residue base-stacking with bound cyclic dinucleotides^13^. These data, along with other identified escaper mutations, suggested that Acb2 nucleotide binding was essential to activation of OptSE. We tested this hypothesis using Acb2^Y8A^, a mutation that was previously shown to disrupt cyclic dinucleotide binding (**Fig. 4c**)^12,13^. Acb2^Y8A^ co-expression with *optSE* also failed to inhibit colony formation. These findings suggest that Acb2 binds nucleotides to activate OptSE signaling.

Given that Acb2 activates the OptSE system and is a phage “sponge” that binds a wide array of nucleotide-derived products, we confirmed that Acb2 can bind the *Kp*OptS product, 2′,3′-c-di-AMP. HPLC experiments showed that incubation of purified T4 Acb2 with *Kp*OptS-derived 2′,3′-c-di-AMP depleted detectable signal (**ED Fig. 5a and b**). When treated with proteinase K, Acb2 was degraded, and 2′,3′-c-di-AMP was once again detected by HPLC, indicating that Acb2 functions as a sponge of 2′,3′-c-di-AMP. As expected, the escaper Acb2^F82C^ showed diminished binding (**ED Fig. 5c**). Consistent with these results, isothermal titration calorimetry (ITC) demonstrated that Acb2 binds 2′,3′-c-di-AMP with an apparent K_d_ of 63.7 nM (**Fig. 4d; ED Fig. 5d-f**).

**Figure 5.**
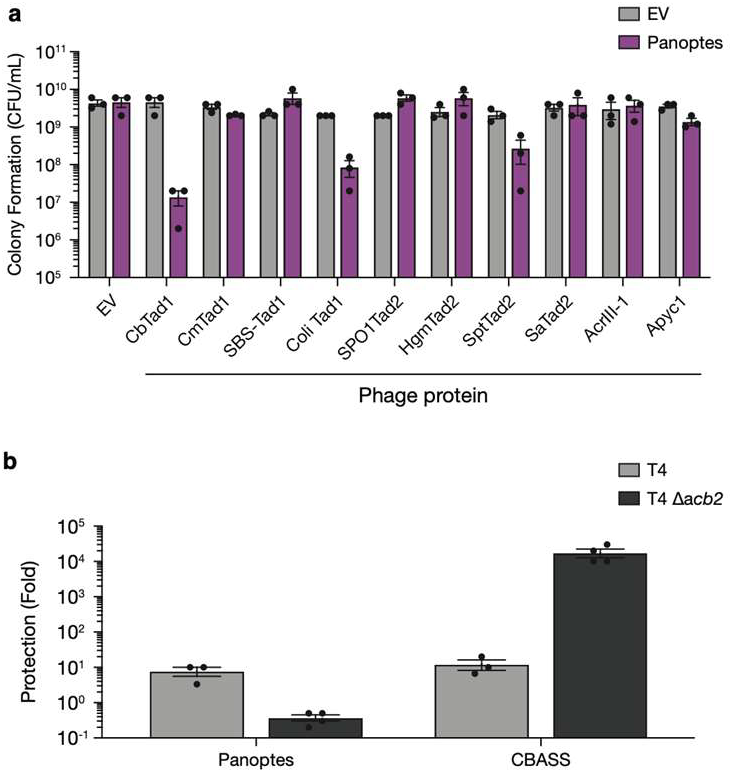
Phage anti-defense proteins that antagonize phage defense activate Panoptes. **(A)** Colony formation of *E. coli* expressing an empty vector or the Panoptes operon on a low copy plasmid and indicated empty vector or phage protein on an IPTG-inducible plasmid. Data represent the mean ± SEM of n = 3 biological replicates, shown as individual points. **(B)** Efficiency of plating of T4 WT or T4 Δ*acb2* phage infecting *E. coli* expressing a plasmid with either an empty vector, Panoptes operon, or CBASS operon. Data represent the mean ± standard error of the mean (SEM) of n = 3 biological replicates, shown as individual points.

Our findings were initially paradoxical. Phage defense systems such as CBASS, Pycsar, and Thoeris each generate nucleotide second messengers to activate defense. OptS and OptE are predicted to generate and respond to cyclic oligonucleotides, respectively. Yet Acb2 nucleotide binding activated, rather than inhibited, OptSE signaling. To explain these observations, we hypothesized that rather than OptS-derived nucleotides *activating* OptE, instead, OptS-derived second messengers *inhibit* OptE. If this were the case, OptE should inhibit growth unless an OptS-derived nucleotide is present.

We tested this hypothesis using plasmids that expressed *optS* and *optE* from IPTG- and arabinose-inducible promoters. Expression of *optS* alone did not impact growth, however, expression of *optE* alone potently inhibited colony formation (**Fig. 4e**). Growth of *optE*-expressing strains could be restored by expressing wild-type *optS* but required the predicted OptS active site residues.

To further confirm our hypothesis, we measured OptS synthesis of 2′,3′-c-di-AMP *in vivo* using a biosensor to interrogate bacterial cell lysates (**ED Fig. 6**). 2′,3′-c-di-AMP could not be detected in *E. coli* expressing empty vector, however, the nucleotide was robustly detected in *optSE*-expressing cells. We estimate the intracellular concentration of 2′,3′-c-di-AMP in *optSE*-expressing cells to be between 165 and 330 nM. These data support a model where, under steady-state conditions, OptS constitutively produces 2′,3′-c-di-AMP to restrain the toxicity of OptE and, thus, depletion of nucleotide by Acb2 unleashes OptE-mediated growth inhibition.

### Diverse phage proteins activate Panoptes

Phages encode a wide variety of genes that allow evasion of bacterial immune pathways that use nucleotide-derived signaling molecules. Thoeris anti-defense proteins 1 (Tad1) and 2 (Tad2) sequester both glycocyclic ADP-Ribose (gcADPR) molecules synthesized by TIR domains during Thoeris defense and cyclic oligonucleotides synthesized by CBASS systems^14,17,24^. Another mechanism that phages use to bypass bacterial immune pathways is through degradation of the nucleotide signaling molecule. Anti-Pycsar 1 (Apyc1), anti-CBASS 1 (Acb1), and type III anti-CRISPR (AcrIII-1) are phage-encoded phosphoesterases that degrade cyclic nucleotide molecules^11,18^. We hypothesized that these phage anti-defense proteins might also activate the OptSE system through either sequestration or degradation of 2′,3′-c-di-AMP molecules synthesized by OptS.

We co-expressed four Tad1 alleles from diverse phages with *optSE* in *E. coli* and observed inhibition of colony formation when either *Cb*Tad1 *or Coli*Tad1 were expressed (**Fig. 5a**). We also tested four different Tad2 proteins and only observed a slight reduction in colony formation when SptTad2 was expressed in the presence of OptSE. These data suggest that other phage “sponge” proteins bind and sequester 2′,3′-c-di-AMP, thereby activating the Panoptes system. On the other hand, we found the phage phosphoesterases Apyc1 or AcrIII-1 did not activate the Panoptes system (**Fig. 5a**), and that *acb1* did not alter Panoptes restriction of T4 phage (**Fig. 4b**). These data indicate that a subset of phage immune evasion proteins can be detected by the Panoptes system, which may be dictated by their nucleotide preference.

### Phyletic patterns and gene-neighborhood linkages of the mCpol-dependent prokaryotic immune systems

mCpol-containing systems are relatively rare but widely distributed across the two prokaryotic superkingdoms (**Extended Data Fig. 7a**). Like the GGDEF and CRISPR polymerase domains, they are strictly excluded from eukaryotes^31^. 53% of all mCpol systems are encoded in genomes also coding for systems centered on SMODS enzymes, a highly significant linkage (p=3.7 × 10^−12^) indicative of functional coupling between them (**Extended Data Fig. 7b and c**). Further, the mCpol genes show a similarly significant coupling with genes coding for members of the S-2TMβ family, like OptE (p=6.9 × 10^−25^). Notably, beyond co-occurring in the same genome as SMODS domain proteins, in roughly half of these cases, *optSE* is also frequently coupled to CBASS systems in the same gene neighborhood as immediately adjacent operons (**Extended Data Fig. 7**). In these instances, the coupled CBASS systems typically encode an effector with either the S-2TMβ or patatin-like phospholipase domain and a Ub-conjugation-like system implicated in the generation of CD-NTase-target protein adducts^13,28,35^. Taken together, these observations strongly suggest that Panoptes systems are predominantly guardians of CBASS systems whose effectors primarily act by compromising the integrity of the cell membrane. Frequently, these coupled systems display two distinct S-2TMβ effectors – one related to Cap15 and one related to OptE (**Extended Data Fig. 7d**) – suggesting an interplay between the versions that are positively and negatively controlled by the nucleotides generated by their respective regulatory synthases. To emphasize the selective benefit or disadvantage of phage T4 expressing *acb2*, we showed that while expressing *acb2* is harmful for phage replication in the presence of the Panoptes defense system, it is required for circumventing a *Vibrio cholerae* CBASS system that synthesizes 3′,3′-cGAMP (**Fig. 5b**).

Beyond coupling with the S-2TMβ, 40% of the mCpol genes are coupled in operons with a gene coding for a protein with a 2TM segment coupled with a CARF domain and a smaller set with a comparable gene encoding a similar 2TM protein with a SAVED domain in lieu of the CARF domain (**Extended Data Fig. 7d**). In these contexts, the CARF and SAVED domains are related Rossmannoid second-messenger sensor domains that take the place of the nucleotide-binding β-barrel domains of the S-2TMβ effectors^36^. Hence, these are predicted to be comparable effectors to the S-2TMβ proteins that are regulated by second-messenger nucleotides. Further, more infrequently, the mCpol genes are coupled in operons or fused to HEPN RNases and an array of predicted membrane perforating 2TM domains. This suggests that the mCpol-generated nucleotides might regulate a wider spectrum of effectors; however, it remains to be seen if they are negative regulators as in the Panoptes system or positive regulators as in the CRISPR and SMODS-based CBASS systems.

## Discussion

Here we discovered that the Panoptes antiphage system flips the liability of a nucleotide-derived second messenger in immune signaling against the phage by detecting the activity of counter-defense proteins. OptS constitutively synthesized 2′,3′-c-di-AMP to hold OptE in an inactive state. During phage infection, Acb2 or similar immune evasion proteins are produced that sequester the OptS-derived signaling molecule, leading to a population of OptE that is no longer bound to cyclic dinucleotide and is free to become activated, in turn aborting infection. While both OptE and Cap15 are closely related members of the S-2TMβ family, they appear to use opposite signaling modalities. OptE inhibited growth in the absence of cyclic nucleotide, whereas Cap15 inhibited growth in the presence of cyclic nucleotide. We anticipate that the growth-inhibiting state of both proteins is the oligomer, which disrupts membrane integrity, but that subtle differences alter whether nucleotide binding stabilizes that monomer or oligomer.

The Panoptes system forces a dilemma for phage: should the phage lose Acb2 and be susceptible to CBASS or evade CBASS and risk agonizing Panoptes. Phage T4 has maintained a wild-type Acb2, however, Phage T2 encodes a mutation that likely selectively disrupts the Acb2 cyclic dinucleotide binding site and the related *Pseudomonas aeruginosa* phage PaMx41 encodes a premature stop codon in *acb2*^12^. Phage can solve this dilemma by tuning the specificity of Acb2 away from the OptS-derived nucleotide, using an alternative CBASS-evasion protein such as Acb1 with different nucleotide specificity, or employing CBASS antagonists that cripple the nucleotide synthase, called a CD-NTase, directly^37^. However, these responses by phage are limited by a key feature of the bacterial immune system: antiphage systems are distributed throughout that bacterial pangenome. In this way, Panoptes and CBASS can be encoded in the same genome so long as they use slightly different nucleotides or as different genes and use the identical nucleotide. The latter circumstance would result in antiphage system incompatibility– an interesting additional layer to the ecology of phage defense gene distribution. Undoubtedly, Panoptes is partially responsible for the evolutionary pressures that have selected for an astonishing diversity of CD-NTase products found in CBASS systems.

Panoptes joins a growing list of antiphage systems that detect immune evasion, such as PARIS and PrrC^38,39^. However, a key difference is that Panoptes surveys the cell using a decoy molecule to sense the activity of anti-defense, as opposed to directly binding the proteins of anti-defense. We speculate that bacteria must encode yet-to-be-discovered OptSE-like systems that guard Pycsar- and Thoeris-derived second messenger pools too. The discovery of these systems has the exciting potential to identify further protein folds capable of synthesizing diverse nucleotide-derived second messengers. In terms of domain architecture and structure, the minimal polymerase catalytic palm domain seen in mCpols is in stark contrast to the incorporation of the cognate domain in complex domain architectures both in Type III CRISPR Cas10 polymerases and GGDEF domains involved in physiological signaling. This is in keeping with the hypothesis that mCpol-centered systems like Panoptes are surviving remnants of ancient antiviral systems that were subsequently elaborated and incorporated as the second messenger signaling arm of a subset of the CRISPR systems^31,36^.

Like bacteria, eukaryotes also guard immune pathways by surveying for immune evasion often in a process called “effector-triggered immunity”^40–42^. Mammals use nucleotide-derived second messengers for immune signaling and mammalian viruses degrade second messengers in the cGAS-STING and OAS-RNaseL pathways using poxin and 2′,5′-phosphodiesterases, respectively^15,16^. It is highly likely that further mechanisms can be used by these viruses to disrupt second messenger pools, including sponge proteins analogous to Acb2. Undiscovered layers of human immune pathways may also exist that detect these viral evasion mechanisms in a manner similar to OptSE defense systems. Intriguingly, there are many SMODS and TIR domain enzymes of unknown function in several metazoans that could mediate comparable activities^43,44^.

The Panoptes system represents a novel reformulation of basic functional units that are used in other antiphage systems. However, subtle molecular changes have enabled a new mechanism for sensing phage infection through detecting immune evasion. Although the phage would appear to be powerless to escape the combination of CBASS and Panoptes, future research will surely uncover the inevitable phage-encoded anti-anti-anti-defense.

## Acknowledgements

The authors would like to thank Annette Erbse and the Shared Instruments Pool (RRID: SCR_018986) of the Department of Biochemistry at the University of Colorado Boulder for providing access to the Avanti JXN 26 Super Speed centrifuges and rotors, which are funded by NIH Grant R24OD033699-01, as well as the ITC 200, funded by the NIH Shared Instrumentation Grant S10RR026516; and members of the Whiteley and Morehouse labs for their advice and helpful discussion. We wish to thank the UCI Mass Spectrometry Facility for assistance with small molecule analyses and accurate mass measurements. Use of the Stanford Synchrotron Radiation Lightsource, SLAC National Accelerator Laboratory, is supported by the U.S. Department of Energy, Office of Science, Office of Basic Energy Sciences under Contract No. DE-AC02-76SF00515. The SSRL Structural Molecular Biology Program is supported by the DOE Office of Biological and Environmental Research, and by the National Institutes of Health, National Institute of General Medical Sciences (P30GM133894). This work was funded by the National Institute of General Medical Sciences of the National Institutes of Health under award number R35GM157311 (B.R.M); the National Institutes of Health through the NIH Director’s New Innovator award DP2AT012346 (A.T.W.); the PEW Charitable Trust Biomedical Scholars Award (A.T.W.); the Boettcher Foundation Webb-Waring Biomedical Research Award (A.T.W.); the Burroughs Wellcome Fund PATH Award (A.T.W.); a University Colorado ABNexus Grant (joint with A.T.W and Kelly Doran); and a Mallinckrodt Foundation Grant (A.T.W.). A.E.S was supported in part by the NIH Interdisciplinary Predoctoral Training in Molecular Biophysics grant (T32GM145437); K.S. was supported in part by the Undergraduate Research Opportunities Program Individual Grant funded by CU Boulder; and C.R.K.H was supported in part by the Boettcher Foundation Collaboration Grant. U.T. was supported as a fellow of the Cancer Research Institute Irvington Postdoctoral Fellowship (CRI4043). A.N. was supported by Graduate Assistance in Areas of National Need (GAANN) Fellowship #P200A240034 through the University of California Irvine Department of Molecular Biology and Biochemistry. D.S.I. was supported in part by funding through a University of California Irvine Undergraduate Research Opportunities Program Fellowship. A.M.B. and L.A. are supported by the intramural funds of the National Library of Medicine, NIH, DHHS (ZIA LM594244).

## Author contributions

Conceptualization, A.E.S., A.T.W., and B.R.M.; Methodology, A.E.S., A.T.W., and B.R.M.; Investigation, A.E.S., A.N., K.S., D.M.D., M.L.D., E.M.Q.E., C.R.K.H., D.S.I., T.A.N., C.M.N., A.M.B., L.A., A.T.W., and B.R.M.; Resources, U.T. and E.M.K.; Writing – Original Draft, A.E.S., A.T.W, and B.R.M.; Writing – Review & Editing, A.E.S., A.T.W, and B.R.M.; Visualization, A.E.S., D.S.I., and B.R.M.; Supervision, A.T.W. and B.R.M; Funding Acquisition, A.T.W. and B.R.M. Note: The order of authors listed 4^th^–11^th^ is alphabetical by last name.

## Data availability

The crystal structure data for OptS have been deposited in the PDB (9MNR). All other data are available in the manuscript or the supplementary materials. Source data are provided.

## Materials and Methods

*Bacterial strains and growth conditions*

*E. coli* strains that were used in this study are listed in **Supplementary Table 1**. All bacterial cultures were grown in 3.5 mL of media in 14 mL culture tubes shaking at 220 rpm at 37 °C, unless otherwise indicated. “Overnight” cultures are defined as those that were grown for 16–20 hours following inoculation from a single colony or glycerol stock. Where applicable, culture media was supplemented with carbenicillin (100 µg/mL) and/or chloramphenicol (20 µg/mL) for plasmid maintenance or strain selection. *E. coli* OmniPir^45^ was used for strain construction and storage of plasmids and *E. coli* MG1655 (CGSC6300) was used for all phage, colony formation, and nucleotide extraction experiments.

All *E. coli* cultures used for cloning, strain construction, protein expression, and indicated colony formation assays were grown in LB medium (1 % tryptone, 0.5

% yeast extract, and 0.5 % NaCl). All strains were frozen for long-term storage in LB plus 30 % glycerol (v/v) at −70 °C. Strains used to perform phage amplification, phage infection assays, 2′,3′-c-diAMP extraction and measurement, and indicated colony formation assays were cultivated in “MMCG” minimal medium (47.8 mM Na_2_HPO_4_, 22 mM KH_2_PO_4_, 18.7 mM NH_4_Cl, 8.6 mM NaCl, 22.2 mM glucose, 2 mM MgSO_4_, 100 mM CaCl_2_, 3 mM thiamine, Trace Metals at 0.1× (Trace Metals mixture T1001, Teknova, final concentration: 8.3 μM FeCl_3_, 2.7 μM CaCl_2_, 1.4 μM MnCl_2_, 1.8 μM ZnSO_4_, 370 nM CoCl_2_, 250 Nm CuCl_2_, 350 nM NiCl_2_, 240 nM Na_2_MoO_4_, 200 nM Na_2_SeO_4_, 200 nM H_3_BO_3_)). When a strain with two plasmids was cultivated in MMCG medium, bacteria were grown in carbenicillin (20 μg/mL) and chloramphenicol (4 μg/mL). When growing strains that required induction, 500 μM IPTG or 0.2 % arabinose was used to induce, as appropriate. MMCG and LB agar plates contain 1.6 % agar and media components described above.

### Plasmid construction

The plasmids used in this study are listed in **Supplementary Table 1**. Cloning and plasmid construction were performed as previously described^6^. Briefly, genes of interest were amplified from phage genomic DNA or previously constructed plasmids using Q5 Hot Start High Fidelity Master Mix (NEB, M0494L), or were synthesized as FragmentGENEs (Genewiz). Gene inserts were flanked by at least 18 base pairs of homology to the vector backbone outside of the restriction digest sites. Ligation of genes into the digested, linearized backbone vector was done using modified Gibson Assembly^46^ with HiFi DNA Assembly Master Mix (NEB, E2621L). Gibson assemblies were transformed by electroporation into competent OmniPir and plated onto LB (1.6 % agar) plates with appropriate antibiotics to select for successful transformations. Phage genes and *OptS* with point mutations were generated by amplifying the gene of interest in two parts from a plasmid template, with the desired mutation occurring in the overlapping region between the two amplicons. Unless otherwise indicated, all enzymes were purchased from New England Biolabs.

For the *OptSE* operon in a pLOCO3 backbone, complete vectors with the indicated operons were generated as ValueGENEs (Genewiz). The pLOCO3 vector (including sfGFP) was initially constructed using Gibson assembly to join and circularize two FragmentGENEs, one with the pLOCO3 backbone, and one with the sfGFP gene (Genewiz; gene fragment sequences listed in **Supplementary Table 1**).

For all vectors using the pTACxc backbone, pAW1608 was amplified and purified from OmniPir. Purified plasmid was then linearized using BamHI-HF and NotI-HF, or EcoRV-HF and PstI-HF. Gibson ligation was used to circularize the plasmid with the new insert.

For all vectors using the pBAD30 backbone, pAW1640 was amplified and purified from OmniPir. Purified plasmid was then linearized using EcoRI-HF and NotI-HF. Gibson ligation was used to circularize the plasmid with the new insert.

For all vectors using the pET16SUMO2 backbone, pAW1123 was amplified and purified from Sure1. Purified plasmid was then linearized using BamHI-HF and NotI-HF. Gibson ligation was used to circularize the plasmid with the new insert.

Plasmid sequences were verified with Sanger sequencing (Quintara Biosciences, Azenta, and/or Plasmidsaurus). Reads were mapped to the predicted plasmid sequence using the Map to Reference feature of Geneious Prime (default settings).

### Phage amplification and storage

The phages used in this study are listed in **Supplementary Table 1**. Information on the BASEL collection used in this study is reported in Maffei et al. 2021^47^. Phage lysates were generated via plate amplification using a modified double agar overlay^48^. For plate amplification, 400 mL of mid-log MG1655 were mixed with

3.5 mL MMCG soft agar mix (MMCG with 0.35 % agar and 10 mM MgCl_2_, 10 mM CaCl_2_, and 100 mM MnCl_2_) and 100-1,000 PFU. Plates were then incubated overnight at 37 °C. Phages were collected by adding 5 mL of SM buffer (100 mM NaCl, 8 mM MgSO_4_, 50 mM Tris-HCl pH 7.5, 0.01 % gelatin) to the plate and incubating for 1 hour at room temperature. To increase phage titers, the top agar overlay was scraped and harvested along with the SM buffer. The SM buffer and top agar mixture was centrifuged at 4,000 × g for 10 minutes and the supernatant was transferred to a new tube. The resulting liquid was passed through a 0.2 mm filter or treated with 2–3 drops of chloroform, followed by vortexing, to remove viable bacteria. All amplified phages were stored at 4 °C in SM buffer.

### Phage infection assays

Phage infection assays and phage titer quantifications were performed using a modified double agar overlay technique^48^. Strains containing the indicated plasmids were cultivated overnight in MMCG medium (including appropriate antibiotics) and were diluted 1:10 in fresh medium the following day. The bacteria were grown until they reached mid-logarithmic phase (OD_600_ 0.1-0.8). 400 µL of mid-log bacteria were mixed with 3.5 ml 0.35 % MMCG agar (plus 5 mM MgCl_2_ and 0.1 mM MnCl_2_) and poured on top of a MMCG (1.6 % agar) plate. The plate was allowed to cool for 15 minutes. Once cooled, 2 μL of a phage 10-fold serial dilution series was spotted onto the soft agar overlay and allowed to dry, after which the plates were incubated at 37 °C overnight. Plates were imaged ∼18-24 hr after infection.

The resulting phage titer was quantified in PFU/mL for each phage lysate tested. PFU were enumerated based on the lowest phage dilution spot with individual, quantifiable PFU. The dilution at that spot was used to calculate the PFU/mL appropriately. When there was a hazy zone of clearance rather than identifiable plaques, the lowest phage concentration at which this was seen was counted as ten plaques. When there was no clearance observed, the least dilute spot was counted as 0.9 plaques, and this was used as the limit of detection for the assay. Phage infection data was reported as PFU/mL ± standard error of the mean (S.E.M) of n = 3 biological replicates.

### Phage infection time course in liquid culture

Bacterial strains containing the indicated plasmid were grown overnight in 25 mL MMCG media plus appropriate antibiotics. Cultures were diluted to an OD_600_ of 0.1 in fresh media without antibiotics (30 mL total volume) and were grown for two hours at 37 °C with shaking at 220 rpm. After two hours, the cultures were infected with phage at the indicated MOI. The OD_600_ of the cultures was measured at the indicated time points after infection. To enumerate PFU at each time point, 250 µL of culture was harvested and centrifuged at 20,000 × g for 5 min at 4 °C. The supernatant was transferred to a new tube and 3–5 drops of chloroform were added, followed by vortexing, to kill any bacteria that remained. The resulting lysates were titered using the phage infection assay protocol explained above.

### Escaper phage generation and amplification

T4 escaper phages were generated from three unrelated, clonal T4 lysates (“parents”) that were separately plate amplified on wild-type *E. coli* MG1655. To make the T4 escaper phages (“daughters”), 400 µL of mid-log bacteria expressing the *OptSE* operon (in MMCG plus 100 µg/mL carbenicillin) was mixed with 100 µL of parent T4 lysate (∼4 × 10^5^ PFU) and 3.5 mL MMCG top agar and poured onto a MMCG agar plate. The plate was allowed to dry and was incubated overnight at 37 °C. The next day, five single escaper plaques were individually isolated from each parent T4 plate using a Pasteur pipette and soaked in 500 µL SM buffer in an Eppendorf tube. A dilution series of each T4 escaper phage was spot plated onto *E. coli* MG1655 expressing the OptSE system to confirm replication in the presence of OptSE. From this plate, single plaques from each escaper were individually purified and plate amplified (as described in the phage amplification and storage protocol above) before storage.

### Phage genome sequencing and escaper analysis

The genomes of the parent and escaper T4 phages were purified as previously described^49^. To do this, 450 mL of phage lysate (>10^8^ PFU/mL) was mixed with 50 µL 10× DNAse I buffer (100 mM Tris-HCl pH 7.6, 25 mM MgCl_2_, and 5 mM CaCl_2_) and treated with DNAse I (final concentration 2 × 10^−3^ U/µL) and RNAse A (final concentration 2 × 10^−2^ mg/mL). This mixture was incubated for 1.5 hours at 37 °C to remove extracellular nucleic acids. After, EDTA was added to a final concentration 20 mM to stop the reaction. Each parent and escaper phage genome was then isolated and purified using the Qiagen DNeasy cleanup kit, starting at the proteinase K digestion step^49^.

The purified phage genomes were prepared for Illumina sequencing using a modification of the Nextera kit protocol as previously described^50^. Illumina sequencing was performed using a MiSeq V2 Micro 300-cycle kit (SeqCenter, LLC). The resulting reads were mapped to Genome accession AF158101.6 using Geneious software’s Map to Reference feature. Each read was trimmed to remove the Nextera adapter sequences before mapping (sequence trimmed: AGATGTGTATAAGAGACAG) using the ‘‘Trim sequences’’ option; otherwise, Geneious default settings were used. The trimmed sequences were mapped to the phage genome using default settings with the “Map to Reference(s)” feature. The Geneious feature “Find Variations/SNPs” was used to identify variants in daughter phage genomes. Called variants were identified as escaper mutations if they were present in ≥75% of reads and were not present in parent phage genomes.

### Construction of phage gene deletions

T4 knockout phages were generated as previously described^51^. Briefly, 5 mL of *E. coli* MG1655 expressing a pET vector encoding the template for homologous repair were grown in LB to mid-log phase. Bacteria were then infected with ∼4 × 10^8^ PFU of T4 and grown for 2–3 hours before harvesting lysate. Phage lysates amplified in this way were then mixed with 400 μL mid-log bacteria expressing eLbuCas13a constructs with spacers targeting the gene of interest and poured onto an MMCG agar plate as in the method described in solid plate amplification detailed above. Individual plaques were isolated and spot-plated on *E. coli* MG1655 expressing the same spacer to confirm mutation and to plaque-purify each clone. Target gene deletion was validated using PCR.

Homologous repair templates encoded 250 bp of homology on either side of the target gene. In-frame deletions retained the first and last 6 amino acids of the target gene, but deleted the intervening sequence. Two 31 nt spacers were selected to target the beginning of each gene and induced as needed using anhydrotetracycline at 5 nM in the top agar of the soft agar overlay.

The *acb1* knockout was constructed with repair template pEK0220 and spacers pEK0223 and pEK0224. The knockout was PCR verified using the primers oEK0490 and oEK0491.

The *acb2* knockout was constructed with repair template pEK0221 and spacers pEK0225 and pEK0226. The knockout was PCR verified using the primers oEK0492 and oEK0493.

The double knockout was created by doing the Δ*acb2* knockout steps on a confirmed Δ*acb1* knockout phage.

### Colony formation assays for measuring bacterial growth inhibition

Bacterial growth inhibition was tested using colony formation assays. Bacterial strains with indicated plasmids were grown overnight in MMCG media plus appropriate antibiotics. The cultures were then 10-fold serially diluted in fresh MMCG media (without antibiotics) and 5 µL of each dilution was spotted onto an MMCG agar plate containing the appropriate antibiotics, as well as IPTG (500 µM; induced condition) as indicated. Data in **Fig. 4e** was collected using LB media with appropriate antibiotics, with or without glucose (0.2% w/v; uninduced condition), IPTG (500 µM; induced condition), or arabinose (0.2% w/v; induced condition), as indicated. After the spotted bacteria was allowed to dry, plates were incubated at 37 °C for ∼16–18 hr for LB plates or ∼24 hr for MMCG plates. Growth inhibition was quantified the next day by counting the number of colony forming units (CFU) of the lowest dilution that had individual colonies. When no individual colonies could be counted, the lowest bacterial concentration at which growth was observed was counted as ten CFU. In instances where no growth was visible, the least dilute spot was counted as 0.9 CFU and used as the limit of detection. Colony formation data was reported as CFU/mL ± S.E.M of n = 3 biological replicates.

### Acb2 protein expression and Isothermal titration calorimetry

The vector expressing WT 6xHis-hSUMO2-Acb2 was transformed into Rosetta2 expressing the pRARE2 plasmid and plated onto 1.6% MMCG agar plates + 100 μg/mL carbenicillin and 20 μg/mL chloramphenicol. An individual colony was picked the following day and inoculated into 100 mL of M9ZB media (47.8 mM Na_2_HPO_4_, 22 mM KH_2_PO_4_, 18.7 mM NH_4_Cl, 85.6 mM NaCl, 1% Casamino acids (VWR), 0.5% v/v Glycerol, 2 mM MgSO_4_, Trace Metals at 0.5 × (Trace Metals Mixture) plus 100 μg/mL carbenicillin and 20 μg/mL chloramphenicol. The culture was then grown overnight shaking at 37 °C and 220 rpm. The following day, the culture was used to inoculate 2 L of the same, fresh media to an OD_600_ of 0.05, then grown to an OD_600_ of ∼1.5. Cultures were crash-cooled on ice for 30 minutes before IPTG was added to 500 μM to induce protein expression. The culture was then moved to a 16 °C shaking incubator and allowed to grow overnight.

Cultures were harvested by centrifugation for 30 minutes at 5,000 rpm and 4 °C in an Avanti JXN-26 Floor Centrifuge using the JXN 12.500 rotor (Beckman). The resulting pellets were resuspended in 40 mL Lysis buffer (20 mM HEPES pH 7.5, 400 mM NaCl, 10% v/v Glycerol, 20 mM Imidazole, 0.1 mM Dithiothreitol (DTT)). After resuspension, cells were lysed by sonication at 80% amplitude, with 15 second on, 45 second off pulses for a total processing time of 10 minutes using a Q500 sonicator (Qsonica). Cellular debris was removed from sonicated lysates by centrifugation for 45 minutes at 4 °C and 16,000 × g in an Avanti JXN-26 Floor Centrifuge using the JA 25.50 rotor (Beckman). The soluble lysate was then decanted and protein was purified using immobilized metal affinity chromatography. Briefly, the soluble lysate was run over 2 mL of HisPure cobalt slurry (Fisher Sci) equilibrated in Lysis Buffer. The resin was then washed with 2 × 25 mL of Wash Buffer (20 mM HEPES pH 7.5, 1 M NaCl, 10% v/v glycerol, 20 mM Imidazole, 0.1 mM DTT) and protein was eluted in 10 mL of Elution Buffer (20 mM HEPES pH 7.5, 400 mM NaCl, 10% v/v glycerol, 300 mM Imidazole, 0.1 mM DTT). Proteins were then dialyzed against 2 × 1 L of Dialysis Buffer (20 mM HEPES pH 7.5, 250 mM KCl, 0.1 mM DTT), overnight at 4 °C using 3.5 kDa MWCO Snakeskin Dialysis Tubing (VWR). The 6×His-SUMO-tag was cleaved using 6×His-hSENP2 (produced in-house; final concentration of 1:100 hSENP2:protein w/w) during the overnight dialysis step. After dialysis, proteins were run over 2 mL HisPure cobalt slurry equilibrated in dialysis buffer to remove any 6×His-SUMO tagged proteins.

After dialysis, the protein was concentrated as needed using 3 kDa MWCO Nanosep spin concentration columns (Pall Labs) and stored 200–500 μL aliquots in Dialysis Buffer at –70 °C. Protein concentrations were measured using A_280_ on a Nanodrop OneC (Thermo) and protein purity was visualized using SDS-PAGE followed by Coomassie staining.

ITC assays were adapted from the protocol described in Huiting et al., 2023. Briefly, the K_d_, ΔH, and ΔS for the binding of WT Acb2 with 2′,3′-c-di-AMP were determined using a MicroCal ITC200 calorimeter. Purified Acb2 and 2′,3′-c-di-AMP were dialyzed into the ITC buffer (20 mM HEPES pH 7.5 and 200 mM NaCl) at 4 °C overnight. The titration was carried out with 27 successive, 1.5 uL injections of 150 µM 2′,3′-c-di-AMP into the sample cell containing 50 µM WT Acb2. Each injection was spaced by 180 seconds and the cell was kept at 25 °C and stirred at 750 rpm. Origin software was used for integration and global curve fitting with a single site binding model.

### AbCap5 protein expression and purification

The vector for expressing 6×His-hSUMO2-AbCap5 was transformed into Rosetta2 expressing the pRARE2 plasmid and plated onto 1.6 % LB agar plates (plus 100 µg/mL carbenicillin and 20 µg/mL chloramphenicol). The plate was allowed to dry and was incubated at 37 °C overnight. The next day, an individual colony was picked to inoculate 25 mL of liquid LB media (plus 100 µg/mL carbenicillin and 20 µg/mL chloramphenicol), which was then grown overnight shaking at 220 rpm and 37 °C. The following day, the overnight culture was diluted 1:100 into fresh media plus antibiotics, then grown to an OD_600_ of ∼0.6 prior to IPTG addition to a final concentration of 500 µM to induce protein expression. Each culture flask was then moved to a 16 °C shaking incubator (220 rpm) and allowed to grow overnight.

The next day, cultures were harvested by centrifugation for 30 minutes at 5,000 rpm and 4 °C in an Avanti JXN-26 Centrifuge using the JXN 12.500 rotor (Beckman Coulter). Each bacterial pellet was resuspended to a total volume of 40 mL lysis buffer (20 mM HEPES pH 7.5, 400 mM NaCl, 10% v/v glycerol, 30 mM imidazole, and 1 mM dithiothreitol (DTT)) plus 1 µL of Pierce Universal Nuclease (Thermo Fisher Sci.). The resuspended bacterial pellets were kept chilled and were lysed by sonication at 80% amplitude, with 30 seconds on, followed by 30 seconds off, for a total processing time of 30 minutes using a Sonicator 4000 (Misonix). The lysed bacteria were centrifuged at 4 °C for 1 hr at 14,000 xg in a 5910 R tabletop centrifuge (Eppendorf) to pellet cellular debris left over from sonication.

The resulting soluble lysate was transferred to a new conical tube and kept on ice. To purify protein, the entire soluble lysate was run over 1 mL of lysis buffer-equilibrated Ni-NTA resin (Thermo Fisher Sci.). The flow through was collected and reapplied to the resin. The resin was then washed with 5 × 25 mL of wash Buffer (20 mM HEPES pH 7.5, 1 M NaCl, 10% v/v glycerol, 30 mM imidazole, and 1 mM DTT). The protein was eluted in 10 mL of elution Buffer (20 mM HEPES pH 7.5, 400 mM NaCl, 10% v/v glycerol, 300 mM imidazole, and 1 mM DTT). The eluted protein was added to 10 kDa MWCO tubing (VWR) and then dialyzed in 1 L of Dialysis Buffer (20 mM HEPES pH 7.5, 250 mM KCl, and 1 mM DTT) for 1 hr at 4 °C. Afterwards, the dialysis tubing and protein were placed into 1 L of fresh Dialysis Buffer and allowed to dialyze overnight at 4 °C. The 6×His-SUMO-tag was cleaved from the N-terminus of AbCap5 using 6×His-hSENP2 (purified in-house; final concentration of 1:100 hSENP2:protein w/w), which was added to the purified protein immediately before it was placed in the dialysis tubing. After dialysis and 6x-His-SUMO cleavage, the purified proteins were applied to 1 mL of dialysis buffer-equilibrated Ni-NTA and the flow through was collected and reapplied to the resin to remove any uncleaved 6×His-SUMO tagged proteins.

The purified AbCap5 was concentrated as needed using 30 kDa MWCO Macrosep spin concentration columns (Pall Labs) and stored in 1 mL aliquots in 50% glycerol v/v at –20 °C. A Nanodrop OneC (Thermo Fisher Sci.) was used to measure A_280_ to determine the protein concentration. Protein purity was determined by using SDS-PAGE followed by Coomassie staining. The AbCap5 protein purified in this way was used in nuclease assays for the measurement of intracellular 2’,3’-c-di-AMP.

### Bacterial lysate preparation and nucleotide extraction

Bacterial strains expressing either the *OptSE* operon or an empty vector were grown overnight in 25 mL MMCG plus carbenicillin (100 µg/mL). The following day, the overnight culture was diluted 1:10 into 100 mL (total volume) of fresh media (without antibiotics) and was allowed to grow to an OD_600_ of ∼0.6–0.7 (noting the specific OD_600_ at harvest). Once the appropriate OD_600_ was reached, the cultures were centrifuged at 4,000 × g for 10 min at 4 °C in a 5910 R tabletop centrifuge (Eppendorf) to pellet the bacteria. The resulting supernatant was discarded, and the bacterial pellet was resuspended in 500 µL of lysis Buffer (10 mM Tris-HCl pH 7.5 and 25 mM KCl) and transferred to a new Eppendorf tube. Hen-lysozyme (0.2 mg/mL final concentration; VWR) and ∼200 µL of zirconium beads were added prior to boiling at 95 °C for 2.5 min, and then vortexed at max speed for 2 min. The samples were boiled and vortexed again and then treated with proteinase K (0.03 mg/mL final concentration; Qiagen) and 1 µL Pierce Universal Nuclease (Thermo Fisher Sci.) for 30 min at 37 °C. After treatment, the samples were boiled and vortexed once again and Triton-X was added to a final concentration of 0.5 % v/v. The samples underwent a final boil and vortex, and were centrifuged at 17,500 × g for 10 min at 4 °C. The resulting lysate/supernatant was transferred to a new Eppendorf tube and kept on ice until ready for nucleotide extraction.

Nucleotides were extracted from bacterial lysates using a modified version of a previously described phenol-chloroform/chloroform extraction protocol^12^. Briefly, 450 µL of phenol-chloroform (Thermo Fisher Sci.) was added to 450 µL of bacterial lysate (prepared above), vortexed for 30 sec, and then centrifuged at 17,500 × g for 45 min at 4 °C. 400 µL of the top aqueous layer was carefully removed and added to 400 µL of fresh phenol-chloroform in a new tube. This mixture was vortexed for 30 seconds and centrifuged at 17,500 × g for 10 min at 4 °C. 350 µL of the resulting top aqueous layer was carefully removed and added to 350 µL of chloroform (VWR) in a new tube. The mixture was vortexed again for 30 sec and centrifuged one final time at 17,500 × g for 10 min at 4 °C. The resulting top aqueous layer was removed and placed in a new tube and for storage at –20 °C. Nucleotide extraction from indicated strains was carried out for a total of n = 3 biological replicates.

### Intracellular 2′,3′-c-di-AMP measurements

Commercially available 2′,3′-c-di-AMP (Enzo) was used to make a standard curve by 2-fold serially diluting the nucleotide in nuclease-free water. A dilution series of nucleotide extracts from the tested bacterial strains were made by 2-fold serially diluting the extracts in nuclease-free water. For each 25 µL reaction, AbCap5 (final concentration 50 nM) was incubated with 500 ng of linear PCR-amplified DNA and 5 µL of one dilution of nucleotide extract, 2′,3′-c-di-AMP, or water in Reaction Buffer (final concentrations: 10 mM Tris-HCl pH 7.5, 25 mM KCl, and 10 mM MgCl_2_). The mixture was incubated for 2 hrs at 37 °C, followed by a 5 min boil at 95 °C. Reactions were mixed with 5 µL of 6× Gel Loading Dye plus SDS (NEB) and 20 µL were loaded and visualized on a 2% (w/v) agarose gel stained with SYBR Safe (Thermo Fisher Sci.).

The concentration of intracellular 2′,3′-c-di-AMP ([n.t.]) was calculated using the following equation:

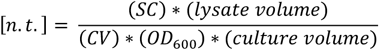

SC is the upper or lower 2′,3′-c-di-AMP standard curve concentration (in [M]) that corresponds to a similar DNA degradation amount as the last visible nucleotide extract dilution reaction (from *OptSE*-expressing cells) that contains non- or partially-degraded DNA PCR product, respectively. Lysate volume is the volume of Lysis Buffer (in L) that was used to resuspend the bacterial pellet before the lysis steps. CV is the OD_600_-specific cell volume (in L * OD^-1^ * mL^-1^) of the bacteria based on the *E. coli* strain and growth conditions^52^. OD_600_ is the optical density of the bacterial culture at the time of harvest (before pelleting). Culture volume is the volume of bacterial culture (in mL) at the time of harvest. Intracellular [2′,3′-c-di-AMP] reported as an average for the upper and lower range of concentrations (in nM) for n = 3 biological replicates of the nucleotide extraction.

### Recombinant protein expression and purification

The genes for full-length *OptS* from *Klebsiella pneumoniae* strain KP67, and bacteriophage T4 Acb2 were codon optimized and synthesized as dsDNA fragments (gBlocks; Integrated DNA Technologies). The genes were cloned using Gibson assembly into a linearized (restriction enzyme digested with BamHI and NotI) in-house pET16 expression vector modified to allow in-frame cloning of an N-terminal hexa-histidine tag with or without a hSUMO2 tag with a Gly-Ser linker. After Sanger sequencing confirmation, the resulting plasmids were subsequently transformed into BL21-RIL *E. coli* cells (BL21 derivative, DE3; Invitrogen, Life Technologies) for protein expression.

Transformant colonies were grown overnight at 37 °C on MDG media (0.5 % glucose, 25 mM Na_2_HPO_4_, 25 mM KH_2_PO_4_, 50 mM NH_4_Cl, 5 mM Na_2_SO_4_, 2 mM MgSO_4_, 0.25 % aspartic acid and trace metals) plates supplemented with 100 μg/mL ampicillin and 34 μg/mL chloramphenicol for selection. Single colonies were used to inoculate liquid MDG media cultures (same components without agar) which were grown overnight ∼16–20 hours at 37 °C and 230 rpm shaking. Overnight cultures were used to inoculate one-liter of liquid M9ZB expression media (0.5 % glycerol, 1 % Cas-amino acids, 47.8 mM Na_2_HPO_4_, 22 mM KH_2_PO_4_, 18.7 mM NH_4_Cl, 85.6 mM NaCl, 2 mM MgSO_4_ and trace metals) contained in 2.5-liter flasks. After cultures reached an OD_600nm_ >2.5, flasks were placed on ice for 15 minutes followed by addition of 0.5 mM IPTG (final) and then incubated at 16 °C with 230 rpm shaking overnight (∼16–20 hours). Cells were harvested by centrifugation, washed with PBS buffer, and flash frozen with liquid N_2_ and stored at −80 °C until needed.

Conventional nickel-affinity chromatography was carried out using gravity flow at 4 °C. Briefly, *E. coli* cell pellets were resuspended with lysis buffer (20 mM HEPES-KOH pH 7.5, 400 mM NaCl, 10% glycerol, 30 mM Imidazole pH 7.5, 1 mM DTT) and subjected to sonication to release cellular contents (10 sec on, 20 sec off, 70% amplitude, 5 min total on time). The lysate was clarified with centrifugation and the supernatant loaded onto 4–6 mL of packed Ni-NTA resin pre-equilibrated with lysis buffer. The column was then sequentially washed with 20 mL of lysis buffer, 70 mL of wash buffer (20 mM HEPES-KOH pH 7.5, 1 M NaCl, 10% glycerol, 30 mM Imidazole pH 7.5, 1 mM DTT), and a final 35 mL of lysis buffer to remove high salt. Protein was eluted with 20 mL of elution buffer (20 mM HEPES-KOH pH 7.5, 400 mM NaCl, 10% glycerol, 300 mM Imidazole pH 7.5, 1 mM DTT). The eluent was dialyzed overnight in 20 mM HEPES-KOH pH 7.5, 250 mM KCl, and 1 mM DTT at 4°C with gentle stirring. In the case of Acb2, hSENP2 protease (D364–L589, M497A) was added to the dialysis tubing to cleave the N6xHis-SUMO2 tag. The dialyzed eluent was concentrated and further purified via size exclusion chromatography (SEC) using a HiLoad™ 16/600 Superdex™ 200 pg column (Cytiva) equilibrated with 20 mM HEPES (pH 7.5), 250 mM KCl, and 1 mM TCEP-KOH. SEC fractions were analyzed with SDS-PAGE and fractions that contained recombinant protein of interest were pooled and concentrated to >10 mg/mL and were flash frozen in liquid N_2_ followed by storage at −80 °C until needed.

### Protein crystallization and structure determination

Apoprotein crystals for OptS from *Klebsiella pneumoniae* strain KP67 were obtained at room temperature by mixing 1 µL of reservoir solution consisting of 8 % (w/v) PEG 4000 with an equal volume of 7 mg/mL of the protein solution using the hanging-drop vapor diffusion method. The crystals were soaked in mother liquor with an additional 15 % ethylene glycol as cryoprotectant and subsequently plunged into liquid N2 and shipped for remote data collection.

X-ray diffraction data for *Kp*OptS were collected at experimental station 12-2 at the Stanford Synchrotron Radiation Lightsource (SSRL) using a Dectris Pilatus® 16M PAD detector. Beamline 12-2 was set at 0.97946 Å and the crystal data was collected at 100 K. The data set was processed using the X-ray Detector Software (XDS) package^53^ that is incorporated into the automated processing pipeline used at SSRL. The structure was determined using molecular replacement conducted by the Phaser-MR program within the PHENIX suite^54^ using a predicted structural model of *Kp*OptS generated by ColabFold v1.5.5 which uses a homology search by MMseqs2 with AlphaFold2^55^. The structure of *Kp*OptS was iteratively refined using the phenix.refine program and the residue positions were manually adjusted in Coot^56^. The final refined structure had the following Ramachandran statistics-98.09 % favored, 1.91% allowed, 0.00% outlier. The full data collection and refinement statistics are summarized in **ED Table 1**. The models were visualized and analyzed using PyMOL (The PyMOL Molecular Graphics System, version 3.1.3, Schrödinger, LLC). The atomic coordinates have been deposited to the Protein Data Bank (PDB ID: 9MNR).

### A*nalysis of enzyme activity with high-performance liquid chromatography*

Enzymatic reactions were performed in 100–600 µL total volume with typically 250 µM total equimolar ribonucleotide triphosphates (ATP, GTP, CTP, UTP), 1 mM MnCl_2_, 10 mM MgSO_4_, 20 mM HEPES-KOH pH 7.4, 100 mM NaCl, and 100 µM OptS protein added last. Reactions were transferred to a 10 kDa-cutoff filter and subjected to centrifugation for 10 minutes at 13,500 x g. When appropriate, purified Acb2was added at 250 µM and incubated for an additional 1 hr at 37 °C. 1 µL of ∼20 mg/mL Proteinase K (NEB, cat no. P8107S) was added to degrade Acb2 and release cyclic dinucleotide before filtering. Likewise, samples were at times treated with P1 nuclease and/or calf-intestinal phosphatase as indicated. All samples were injected at 10 µL onto an Agilent 1200 Series HPLC equipped with a 4.6 × 150 mm and 5 µm particle-size Zorbax Bonus-RP C18 column using an isocratic elution method including 97% 50 mM NaH_2_PO_4_ pH 6.8 and 3% acetonitrile buffer system held at 40 °C. Nucleotide separation was monitored at 254 nm using a multiwavelength detector. For mass spectrometry analysis, enzymatic reactions were prepared using 20 mM ammonium acetate pH 8.0 instead of HEPES-KOH and NaCl but otherwise treated the same with or without an LC fractionation step to isolate specific peaks. For fractionation on HPLC, temperature was kept at 23 °C with a gradient elution method as follows: Solvent A-20 mM ammonium acetate pH 8.0; Solvent B-Methanol. 0–2 min 100% A, 2–8 min 0–100% B, 8–10 min 100% B, 10–11 min 0–100% A, 11–17 min 100% A. ∼1 mL fractions were collected and concentrated via speed-vac (1– 3 hr, 30 °C) prior to injection on a Waters Acquity UPLC-QDA instrument equipped with a C18 50 mm UPLC column and single quadrupole MS detector (QDA; m/z 50-1250) and photodiode array detector (PDA; 200-500 nm). Standard gradient elution was used from 0–100% 0.1 % formic acid:acetonitrile over 5 minutes. In ESI+ mode, analyte m/z ions (M+H)+ or (M+NH3)+ were validated to less than ± 5 ppm relative to the nearest sodiated polyethylene glycol (CAS: 25322-68-3, av. Mwt 400)) or sodiated methoxypolyethyleneglycol (CAS: 990-74-4, av. Mwt 350) calibrant peak lockmass.

### Synthetic nucleotide ligands

Synthetic cyclic dinucleotide ligands used for HPLC and mass spectrometry analysis were purchased from Biolog Life Science Institute: 3′,3′-c-di-AMP (cat no. C 088), 2′,3′-c-di-AMP (cat no. C 187), 2′,2′-c-di-AMP (cat no. C 188), 3′,3′-c-di-GMP (cat no. C 057), 2′,3′-c-di-GMP (cat no. C 182), 2′,2′-c-di-GMP (cat no. C 162), 3′,3′-cGAMP (cat no. C 117), 2′,3′-cGAMP (cat no. C 161), 3′,2′-cGAMP (cat no. C 238), 2′,2′-cGAMP (cat no. C 210), pppA(2′,5′)pA (cat no. T 073).

### Accession numbers

The structure of *Kp*OptS has been deposited in the Protein Data Bank (PDB ID: 9MNR). All other relevant accession numbers can be found in **Supplementary Table 1C**.

### Statistics and reproducibility

Each experiment presented was performed in independent biological duplicate or triplicate using bacterial cultures grown on two or three separate days, respectively. Data was plotted using Graphpad Prism 9 at an n = 2 or n = 3 with error bars indicating the standard error of the mean. Illumina sequencing results were analyzed using Geneious Prime Software.

## Extended Data Tables

Extended Data Table 1. X-ray crystallography data collection and refinement statistics

Extended Data Table 2. T4 phage escaper mutations

## Supplementary Information

Supplementary Table 1. Bacterial strains, plasmids, phages, and proteins used in this study

## Extended Data Figures

**Extended Data Figure 1.**
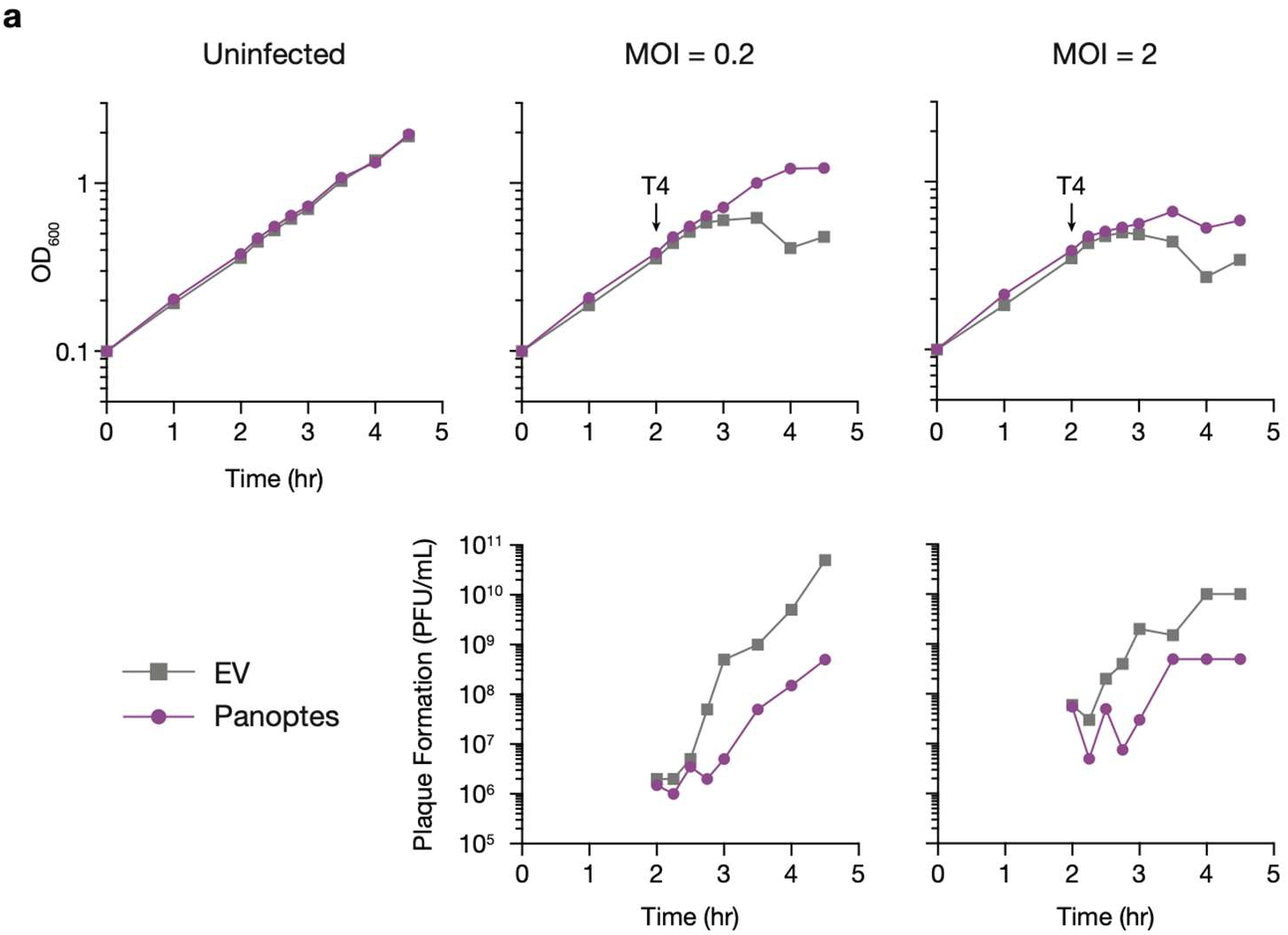
Panoptes protects against phage in liquid culture. **(A**) Above: growth curve of *E. coli* expressing the indicated plasmid. Arrows indicate the time each culture was infected with phage T4 at the indicated multiplicity of infection (MOI). Below: efficiency of plating of the phage present in each sample at the indicated time points. Data are representative of n = 2 biological replicates.

**Extended Data Figure 2.**
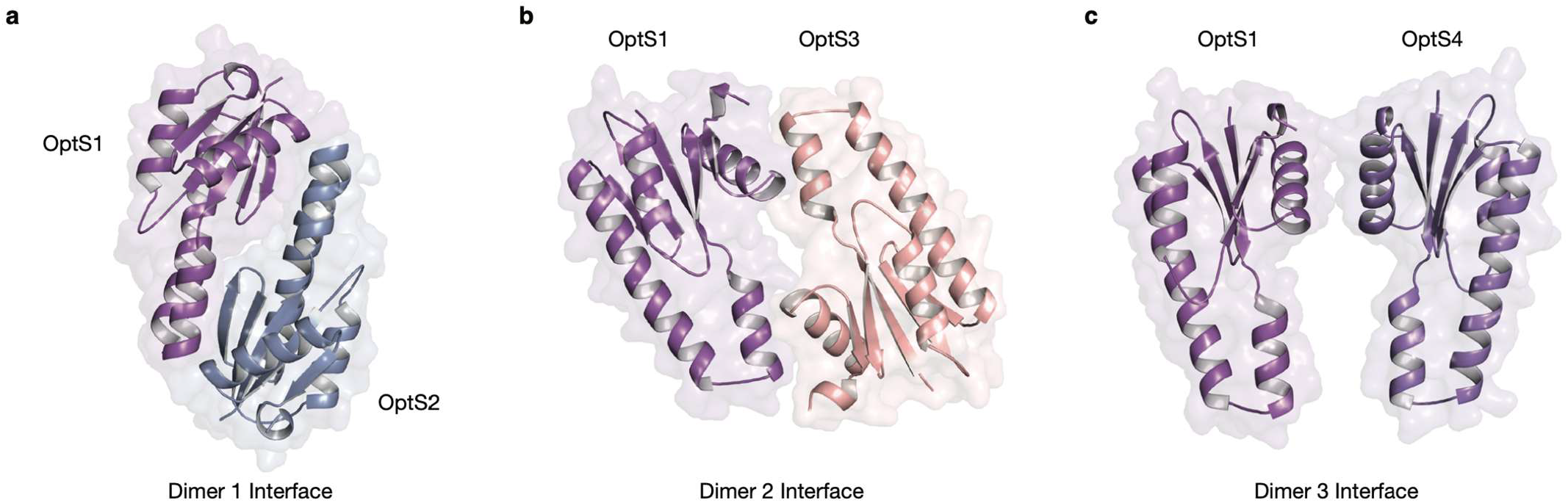
The *Kp*OptS tetramer contains three dimers (pair of protomers) interfaces. **(A)** Dimer 1 interface. **(B)** Dimer 2 interface. **(C)** Dimer 3 interface.

**Extended Data Figure 3.**
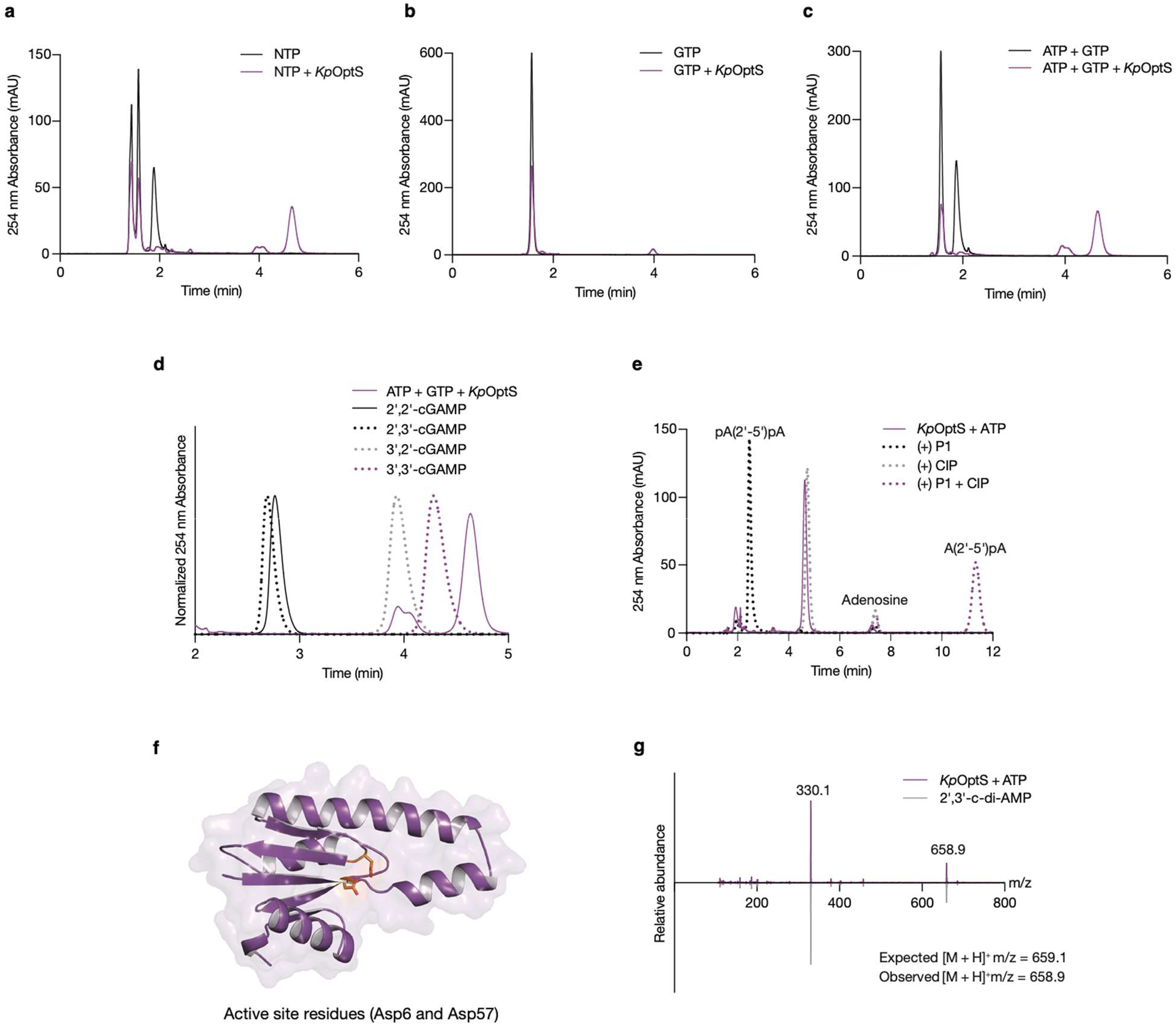
Characterization of *Kp*OptS enzymatic activity *in vitro*. **(A)** HPLC analysis of NTP chemical standards compared with the product of *Kp*OptS when incubated with a mixture of NTPs. Traces representative of at least three replicates. **(B)** HPLC analysis of a GTP chemical standard compared with the product of *Kp*OptS when incubated with GTP alone. Traces representative of at least three replicates. **(C)** HPLC analysis of ATP and GTP chemical standards compared with the product of *Kp*OptS when incubated with ATP and GTP. Traces representative of at least three replicates. **(D)** HPLC analysis of 2′,2′-cGAMP, 2′,3′-cGAMP, 3′,2′-cGAMP, and 3′,3′-cGAMP chemical standards compared with the product of *Kp*OptS. Traces representative of at least three replicates. **(E)** P1 nuclease and CIP treatment of *Kp*OptS ATP-only product. Traces representative of at least three replicates. **(F)** A less magnified view of the putative catalytic site highlighting catalytic Asp6 and Asp57 (orange). **(G)** MS spectra of a 2′,3′-c-di-AMP standard (bottom) and the *Kp*OptS product (top).

**Extended Data Figure 4.**
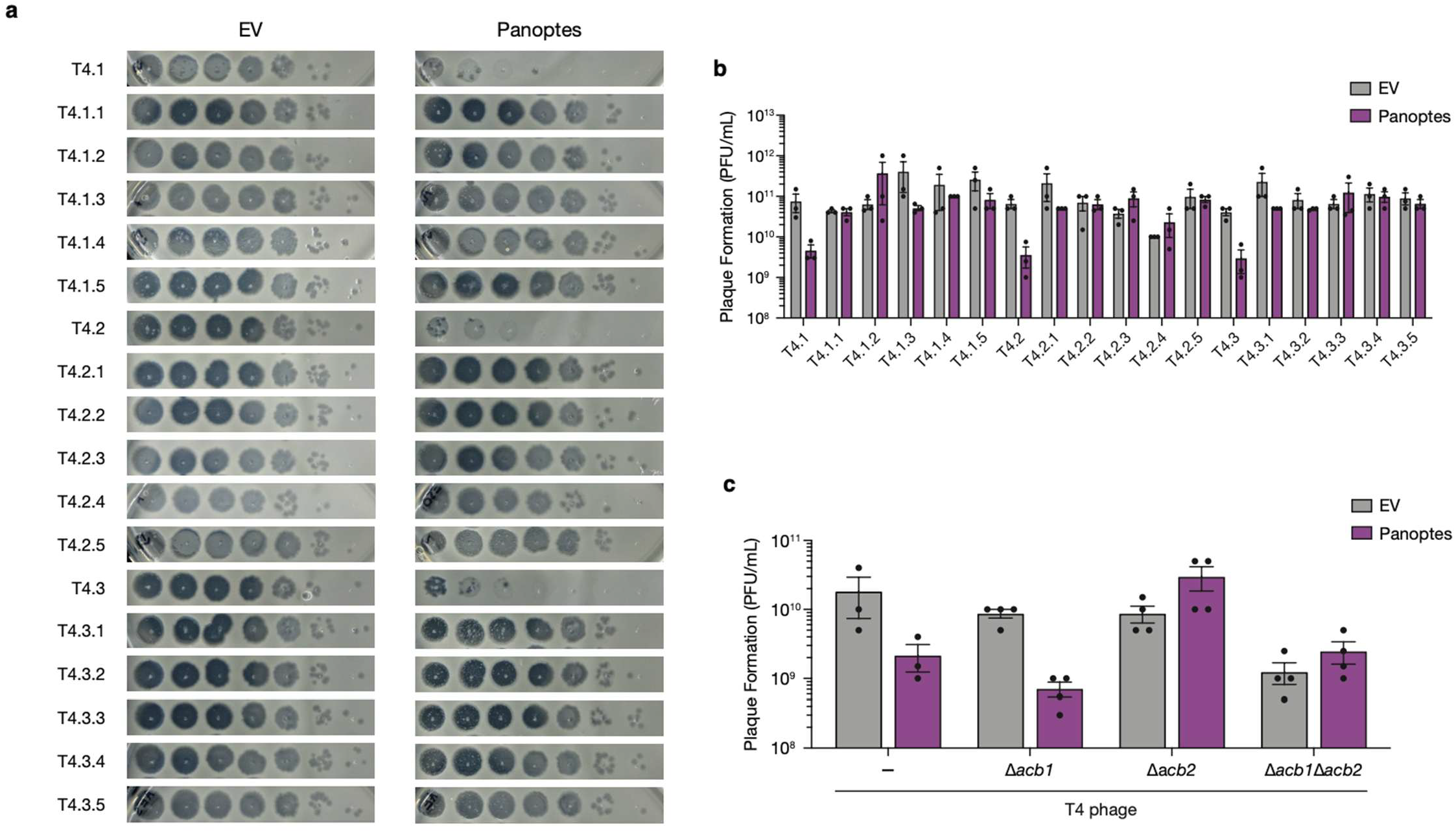
Phage T4 escapers evade Panoptes-mediated defense. **(A)** Efficiency of plating of T4 parent and escaper phages on *E. coli* expressing an empty vector or the Panoptes operon from a plasmid. Images are representative of n=3 biological replicates. **(B)** Efficiency of plating of T4 parent phages and corresponding escaper mutant phages (Escaper, T4.X.Y, where X indicates the parent phage number and Y indicates the escaper phage number) on *E. coli* expressing an empty vector or the Panoptes operon from a plasmid. Data represent the mean ± standard error of the mean (SEM) of n = 3 biological replicates, shown as individual points. **(C)** Efficiency of plating of T4 phages with the indicated genotype on *E. coli* expressing an empty vector or the Panoptes operon from a plasmid. Data represent the mean ± standard error of the mean (SEM) of n = 3 biological replicates, shown as individual points.

**Extended Data Figure 5.**
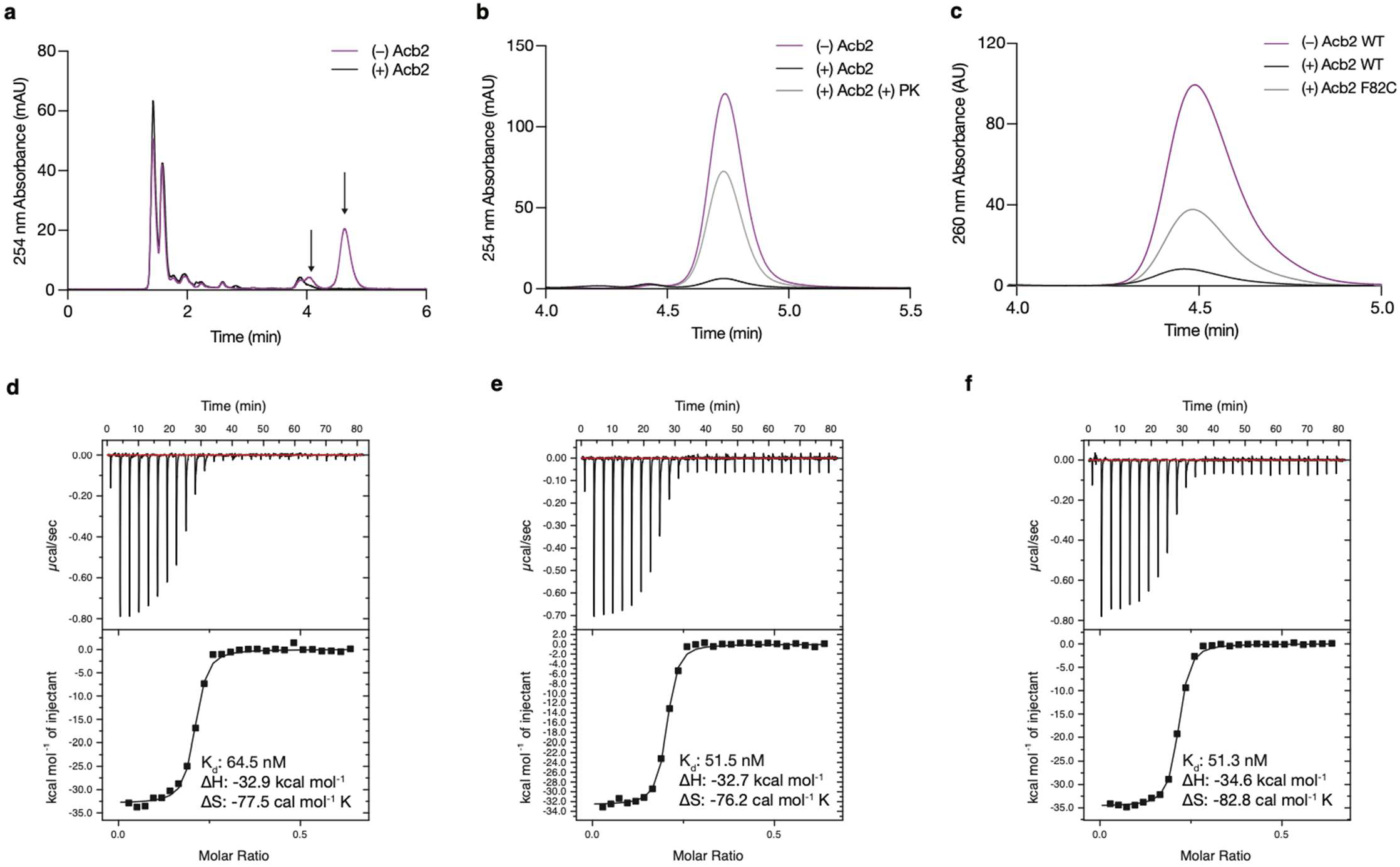
Acb2 binds the product of *Kp*OptS. **(A)** HPLC analysis of the ability of Acb2 to bind the main products of *Kp*OptS (ATP and GTP reaction). Traces representative of at least three replicates. **(B)** HPLC analysis of the ability of Acb2 to bind and release the product of *Kp*OptS when treated with proteinase K (ATP only reaction). Traces representative of >3 replicates. **(C)** HPLC analysis of the ability of mutant Acb2 to bind the ATP-derived product of *Kp*OptS. Traces representative of at least three replicates. **(D-F)** Replicates of ITC assays to test binding of 2′,3′-c-di-AMP to Acb2.

**Extended Data Figure 6.**
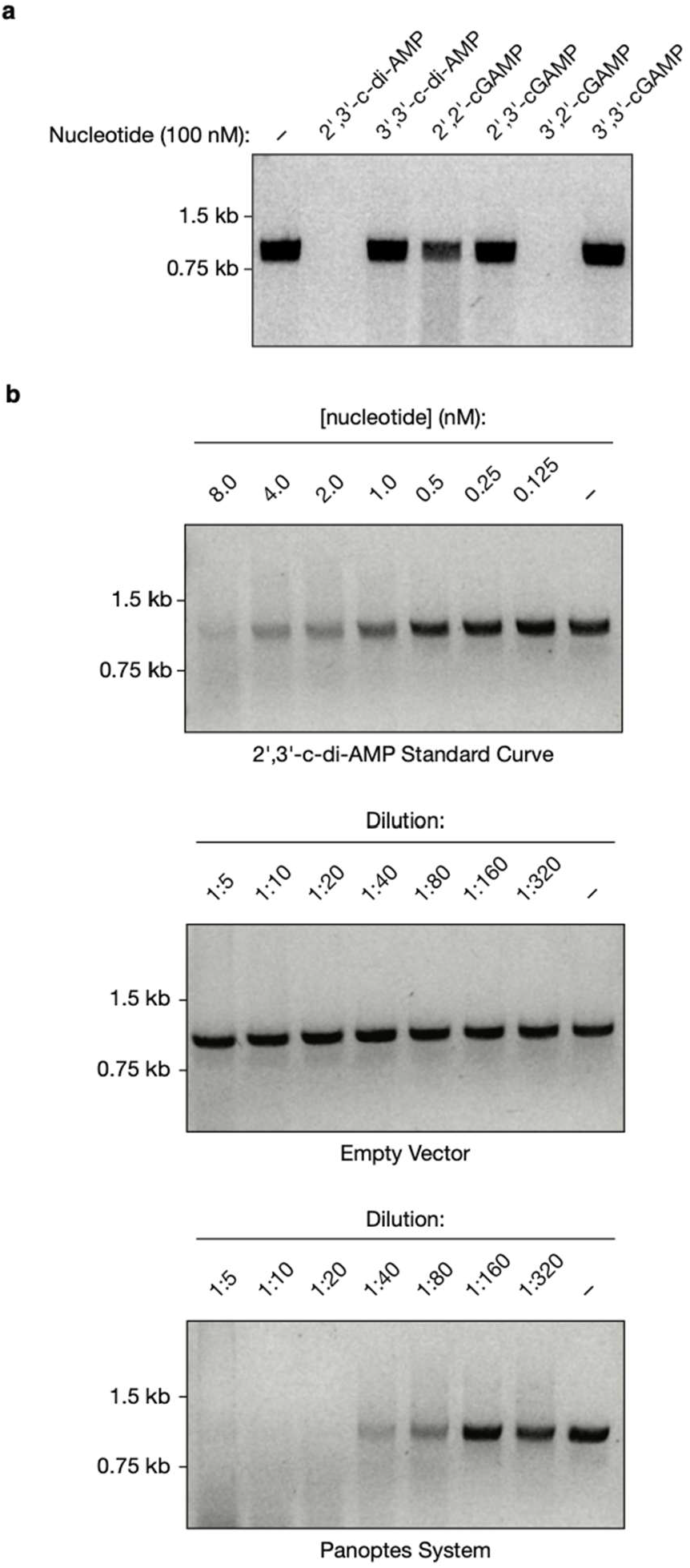
*Ab*Cap5 is a sensor of 2′,3′-c-di-AMP. **(A)** Visualization of DNA degradation by *Ab*Cap5 when incubated with either 2′,3′-c-di-AMP, 3′,3′-c-di-AMP, 2′,2′-cGAMP, 2′,3′-cGAMP, 3′,2′-cGAMP, or 3′,3′-cGAMP. Data are representative images of n=2 technical replicates. **(B)** Visualization of DNA degradation by *Ab*Cap5 when incubated with a dilution series of 2′,3′-c-di-AMP or extracted nucleotides from *E. coli* expressing an empty vector or the Panoptes operon. Data are representative images of n=3 biological replicates.

**Extended Data Figure 7.**
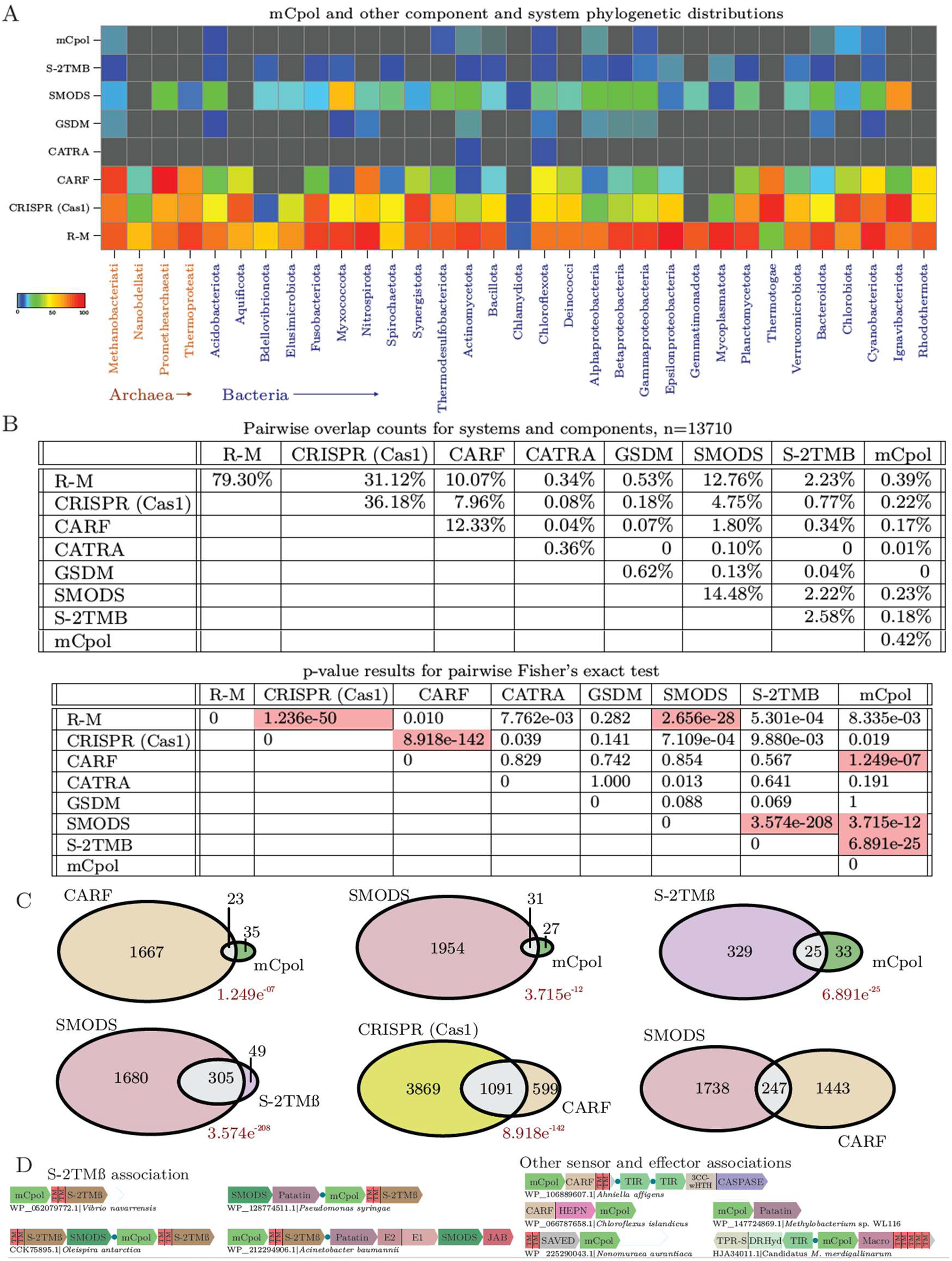
mCpol distribution and gene co-occurrences. **(A)** Heat map depicting genome presence/absence of selected conflict systems in prokaryotic lineages. **(B)** Tables depicting percentages of genome occupancy and pairwise co-occupancy for selected systems across prokaryotes (top) and p-values representing the significance of pairwise co-occurrences from exact Fisher’s tests (bottom). Co-occurrences with p-values of less than 1e-05 are shaded in red. **(C)** Venn diagram depictions of genome overlap between selected pairs, significant p-values are provided below the diagram where applicable. **(D)** Conserved mCpol gene neighborhood associations, with individual genes depicted as boxed arrows. “Core” mCpol-containing systems are separated from discrete associating systems by blue circles.

