## Extended Data Table 1 for "A minimal CRISPR polymerase produces decoy cyclic nucleotides to detect phage anti-defense proteins"

**Table 1 Data collection and refinement statistics (molecular replacement)**

| | *Kp*OptS- apo |
|---|---|
| **Data collection** | |
| Space group | C 1 2 1 |
| Cell dimensions | |
| *a*, *b*, *c* (Å) | 114.17, 39.65, 109.20 |
| α, β, γ (°) | 90.00, 91.99, 90.00 |
| Resolution (Å) | 37.45–1.75 (1.79–1.75)* |
| $R_{merge}$ | 0.066 (0.905) |
| $I / \sigma I$ | 15.5 (1.92) |
| Completeness (%) | 98.3 (96.5) |
| Redundancy | 6.8 (6.2) |
| **Refinement** | |
| Resolution (Å) | 1.75 |
| No. reflections | 48,188 (3414) |
| $R_{work}$ / $R_{free}$ | 0.2214 / 0.2597 |
| No. atoms | |
| Protein | 3702 |
| Ligand/ion | 0 |
| Water | 194 |
| *B*-factors | 33.14 overall |
| Protein | 32.82 |
| Ligand/ion | 0 |
| Water | 39.30 |
| R.m.s. deviations | |
| Bond lengths (Å) | 0.005 |
| Bond angles (°) | 0.66 |

*Data collected from a single protein crystal *Values in parentheses are for highest-resolution shell.
